# Phylogenomics reveals an extensive history of genome duplication in diatoms (Bacillariophyta)

**DOI:** 10.1101/181115

**Authors:** Matthew Parks, Teofil Nakov, Elizabeth Ruck, Norman J. Wickett, Andrew J. Alverson

**Author notes:** these authors contributed equally to this work (Matthew Parks, Teofil Nakov).

## Abstract

**Premise of the study:** Diatoms are one of the most species-rich lineages of microbial eukaryotes. Similarities in clade age, species richness, and contributions to primary production motivate comparisons to flowering plants, whose genomes have been inordinately shaped by whole genome duplication (WGD). These events that have been linked to speciation and increased rates of lineage diversification, identifying WGDs as a principal driver of angiosperm evolution. We synthesized a relatively large but scattered body of evidence that, taken together, suggests that polyploidy may be common in diatoms.

**Methods:** We used data from gene counts, gene trees, and patterns of synonymous divergence to carry out the first large-scale phylogenomic analysis of genome-scale duplication histories for a phylogenetically diverse set of 37 diatom taxa.

**Key results:** Several methods identified WGD events of varying age across diatoms, though determining the exact number and placement of events and, more broadly, inferences of WGD at all, were greatly impacted by gene-tree uncertainty. Gene-tree reconciliations supported allopolyploidy as the predominant mode of polyploid formation, with particularly strong evidence for ancient allopolyploid events in the thalassiosiroid and pennate diatom clades.

**Conclusions:** Whole genome duplication appears to have been an important driver of genome evolution in diatoms. Denser taxon sampling will better pinpoint the timing of WGDs and likely reveal many more of them. We outline potential challenges in reconstructing paleopolyploid events in diatoms that, together with these results, offer a framework for understanding the evolutionary roles of genome duplication in a group that likely harbors substantial genomic diversity.

## INTRODUCTION

Duplicated genes are a hallmark of eukaryotic genomes. For example, some two-thirds of the genes in *Arabidopsis* are present in more than one copy (Ambrosino et al., 2016), a proportion that is typical of most plant genomes (Panchy, Lehti-Shiu, and Shiu, 2016). These duplicated genes can provide raw materials for evolutionary innovation and change, thereby representing an important source of novel traits in lineages spanning the eukaryotic tree of life (Ohno, 1970). In flowering plants, for example, gene duplications have been linked to changes in a diverse set of traits, including floral pigmentation and structure, flowering time, disease and herbivore resistance, fruit characteristics, and stress response (reviewed by Panchy, Lehti-Shiu, and Shiu, 2016). Gene duplication can occur across multiple scales: from small tandem duplications affecting one or a few genes, to transposon-mediated segmental duplications affecting large stretches of a chromosome, to, most dramatically, doubling of the entire genome [whole genome duplication (WGD) or polyploidy] (Flagel and Wendel, 2009; Panchy, Lehti-Shiu, and Shiu, 2016).

The evolutionary history of angiosperms is replete with ancient polyploidy events, such that a majority of the duplicated genes in *Arabidopsis* can be traced to a series of at least four separate WGDs dating back to the origin of flowering plants (Bowers et al., 2003; Jiao et al., 2011). In addition to providing a source of novel and potentially adaptive traits, gene and genome duplications can also serve as mechanisms of speciation (Winge, 1917; Lynch and Force, 2000). Whole-genome duplications, in particular, frequently coincide with speciation events in flowering plants (Otto and Whitton, 2000; Wood et al., 2009; Zhan et al., 2016). An association between WGD and subsequent increases in net diversification rate is also emerging (Otto and Whitton, 2000; Soltis et al., 2009; Tank et al., 2015; but see Kellogg, 2016), implicating WGD as a potentially important driver of species diversification in angiosperms. Polyploidy events have been an important source of novelty in other species-rich lineages as well, including vertebrates (Ohno, 1970; Dehal and Boore, 2005) and fungi (Wolfe and Shields, 1997; Albertin and Marullo, 2012). With longstanding genetic model systems and a wealth of genomic data, these groups are, however, some of the most intensively studied eukaryotes. Growing genomic resources for equally diverse but historically understudied groups have made it possible to explore whether WGD has played a similarly important role in non-model lineages.

With diversity estimates in the tens to hundreds of thousands of species (Guiry, 2012; Mann and Vanormelingen, 2013), a prominent role in the global cycling of carbon and oxygen (Field et al., 1998), a critical position at the base of their native food webs, and a crown age of roughly 200 My (Nakov, Beaulieu, and Alverson, 2017), diatoms are in many respects the angiosperms of the sea. They exhibit many layers of diversity beyond their species richness, including a broad range of ecological niches, life history strategies, and most famously in the diverse patterns and ornamentations of their silicified cell walls (Round, Crawford, and Mann, 1990). Very little is known, however, about the primary sources of genetic change underlying the origins and evolutionary shifts in these traits. Many independent lines of direct and indirect evidence collected over decades suggest that WGD may be common in diatoms. For example, although karyotypes are available for very few species, chromosome counts range from 2n = 8– 130 among raphid pennate species alone (Kociolek and Stoermer, 1989). Flow cytometric measurements have shown substantial variation in genome size, with estimates spanning more than three orders of magnitude among the few dozen species that have been surveyed (Connolly et al., 2008; von Dassow et al., 2008). Within species, a recent genome doubling distinguishes natural populations of the polar centric species, *Ditylum brightwellii* (Koester et al., 2010), and WGDs apparently occur in strains maintained in long-term cell culture as well (von Dassow et al., 2008). Finally, and perhaps most compellingly, simultaneous fusions of three or four gametes, leading to the formation of autopolyploid auxospores (i.e., zygotes), have been directly observed in several raphid pennate diatoms, including *Cocconeis* (Geitler, 1927), *Craticula* (Mann and Stickle, 1991), *Dickea* (Mann, 1994), *Achnanthes* (Chepurnov and Roschin, 1995), and *Seminavis* (Chepurnov et al., 2002). The latter set of observations, in particular, led to the prediction that polyploidy might be an important driver of speciation in diatoms (Mann, 1994, 1999a). Finally, there is some evidence for polyploidy in non-diatom stramenopiles, the higher-order lineage to which diatoms belong (Coyer et al., 2006; Ioos et al., 2006). In light of this relatively large body of evidence, the most surprising discovery might be the lack of a genomic signature for paleopolyploidy in diatoms.

We compiled new and previously sequenced genomic and transcriptomic data for 37 phylogenetically diverse diatom species to estimate, for the first time, the extent to which diatom genomes have been shaped, if at all, by WGD events. Gene counts, gene trees, and patterns of synonymous sequence divergence (Ks) between gene duplicates identified numerous putatively allopolyploid-driven WGDs across the phylogeny and potentially dating as far as back as 200 Mya. We discuss possible modes of polyploid formation in diatoms as well as identifying research directions that will shed light on the mechanisms and evolutionary consequences of WGD in diatoms.

## MATERIALS AND METHODS

### Taxon sampling

We sampled 37 diatom species that spanned the known breadth of extant phylogenetic diversity, the bolidophyte *Triparma pacifica*, and two pelagophyte outgroups (Appendix 1).

### Transcriptome sequencing

We extracted total RNA from exponentially growing cultures using the Qiagen RNeasy kit. We prepared indexed sequencing libraries using the Illumina TruSeq RNA Sample Preparation Kit v2 (Appendix 1). Multiplexed libraries were sequenced with the Illumina HiSeq 2000 or HiSeq 4000 platforms (Appendix 1). Newly generated data were deposited in the Sequence Read Archive databased maintained by the National Center for Biotechnology Information (NCBI) under accessions XXXXXXX–XXXXXXX (Appendix 1 [note: GenBank submissions are pending]).

### Transcriptome assembly and annotation

RNA-seq reads were filtered and assembled following the basic guidelines outlined in the Oyster River Protocol (MacManes, 2015). In short, raw sequence reads were corrected with Rcorrector (Song and Florea, 2015) and quality-trimmed with Trimmomatic (ver. 0.32) (Bolger, Lohse, and Usadel, 2014). Corrected and trimmed reads were filtered for common laboratory vectors and diatom rRNA genes using bowtie2 (ver. 2.2.3) (Langmead and Salzberg, 2012). Overlapping forward and reverse pairs of filtered reads were merged with BBMerge (ver. 8.8) (Bushnell 2014), and both merged and unmerged reads were assembled with Trinity (ver. 2.2.0) (Grabherr et al., 2011b). Assembled transcripts were translated into amino acid sequences using TransDecoder (ver. 2.0.1) (https://transdecoder.github.io/), with translation predictions enabled by BLASTP searches of the longest identified open reading frames to the Swiss-Prot database and HMMER searches (Eddy, 2011) to the Pfam database (Finn et al., 2015). Assembly quality was measured by TransRate scoring (ver. 1.01) (Smith-Unna et al., 2016) and recovery of conserved eukaryotic orthologs present in the BUSCO database (Simão et al., 2015).

### Orthology/Paralogy-based transcriptome clustering

We used CD_HIT (-c 0.99 -n 5) (Fu et al., 2012)to remove redundant isoform transcripts from the full set of amino acid sequences for each species. The non-redundant transcriptome of each species was then searched against a database of all 40 (non-redundant) transcriptomes with BLASTP (ver. 2.3.0+) (e-value ≤ 10^-5^ and max-target sequences = 100) (Camacho et al., 2009), and this output was used to identify putative orthologous clusters with MCL (ver. 12-135) (Van Dongen, 2001; Enright, Van Dongen, and Ouzounis, 2002; Van Dongen and Abreu-Goodger, 2012) with e-value cutoff of 10^-30^ and an inflation value of 1.4. MCL clusters with fewer than four taxa were excluded from subsequent analyses.

### Homolog and species tree reconstructions

Initial orthologous clusters were pruned and resulting ortholog trees were constructed using the ‘phylogenomic_dataset_construction’ pipeline of Yang and Smith (2014). For this pipeline, we aligned sequences with MAFFT (ver. 7.309) (Katoh and Standley, 2013) and reconstructed gene and ortholog trees with RAxML (ver. 8.2.9) (Stamatakis, 2014) using the PROTCATWAG model and 100 rapid bootstrap pseudoreplicates per alignment. As part of the pruning pipeline to create single-copy orthologous clusters for phylogenetic analyses, alignments were trimmed to include only sites with a minimal column occupancy of 0.1, terminal branches with lengths greater than two branch-length units or with lengths greater than 10 times the length of a sister branch were removed, internal branches with lengths greater than 2 branch-length units were removed, and sister tips belonging to the same taxon were reduced to include only the tip with the largest number of unambiguous characters in the trimmed alignment. Yang and Smith’s (2014) ‘RT’ strategy was used to create final, single-copy ortholog alignments, with the two pelagophyte samples specified as outgroup taxa and all diatom samples and *Triparma pacifica* specified as ingroup taxa; this allowed the final set of gene trees to be rooted with a non-diatom outgroup. Finally, we used SumTrees (Sukumaran and Holder, 2010)to collapse nodes on the final ortholog trees with less than 33% bootstrap support.

For species tree reconstructions, ortholog alignments and trees were filtered again to include only those alignments with 100% taxon occupancy and alignment columns with less than 20% missing data or gap characters. Species trees were then reconstructed using both summary-coalescent and concatenation-based approaches. We used ASTRAL (ver. 4.10.8) for summary-coalescent species tree reconstruction, with topology and support estimated with local posterior probabilities (Sayyari and Mirarab, 2016) and multilocus bootstrapping (Seo, 2008). We refer to these as ASTRAL and ASTRAL-mlbs, respectively. For the concatenation-based analysis, models of protein evolution were first determined for each ortholog alignment using ProtTest (ver. 3.4.2) based on the AICc selection criterion (Guindon et al., 2010; Darriba et al., 2011). Alignments were concatenated with AMAS (Borowiec, 2016), and the resulting species tree was inferred using IQ-TREE with ultrafast bootstrapping and SH-aLRT testing (1000 replicates each) (Guindon et al., 2010; Minh, Nguyen, and von Haeseler, 2013; Chernomor, von Haeseler, and Minh, 2016). This tree was used in subsequent analyses as a reference species tree, since we recovered both relatively high levels of gene tree discordance and low levels of gene tree support across input gene trees (see results), under which conditions concatenation-based methods may outperform summary coalescent methods (Mirarab and Warnow, 2015), and its topology was nearly identical to recovered ASTRAL species tree topologies (see results). Gene tree support for this recovered species tree was estimated with PhyParts (analysis=fullconcon) (Smith et al., 2015) and a companion script, phypartspiecharts.py (https://github.com/mossmatters/phyloscripts/tree/master/phypartspiecharts), with gene tree concordance estimated against the IQ-TREE species tree and using a 33% bootstrap support threshold.

The IQ-TREE species tree was time-calibrated using TreePL (Smith and O’Meara, 2012) with 10 fossil-derived calibration points. The minimum and maximum bounds were set following Nakov et al. (2017), except the calibration for the most recent common ancestor of diatoms and Parmales was constrained to a maximum age of 250 million years before present. The optimal rate-smoothing parameter for TreePL was estimated with random-subsample-and-replace cross-validation with a range of tested values on a log scale between 10^5^ and 10^-5^.

### Overall approach to identification of paleopolyploidy events

Identifying WGD events from transcriptome data necessarily relies on temporal or phylogenetic signal, rather than spatial syntenic signal, and so may be impacted by historical variation in molecular evolutionary rates and saturation artifacts (McKain et al., 2016). Nevertheless, several complementary methods are now available that together provide increased confidence in transcriptome-based WGD inferences in the absence of synteny information. These approaches are broadly divided into three categories: (1) paralog divergence (i.e., Ks-based) methods (Lynch and Conery, 2000; Blanc and Wolfe, 2004); (2) gene-tree/species-tree reconciliation methods (Durand, Halldorsson, and Vernot, 2006; Jiao et al., 2011; Thomas, Ather, and Hahn, 2017); and (3) gene count methods (Rabier, Ta, and Ane, 2014). Each of the three approaches provides incrementally more rigorous and specific tests for WGD: (1) the Ks analyses provide semi-quantitative evidence for the presence of synchronously duplicated genes, (2) gene-tree reconciliation pipelines identify specific branches on the species tree with elevated numbers of gene duplications and losses (Durand, Halldorsson, and Vernot, 2006; Yang et al., 2015), a reconciliation approach allows specific tests about the mechanism of WGD events (auto-vs. allopolyploidy) (Thomas, Ather, and Hahn, 2017), and (3) a gene-count method for detecting and locating WGD events independent of both Ks and gene-tree information. As described in the following sections, we applied each of these methods to one or more sets of orthologous clusters and their corresponding gene trees (Fig. 1).

**Figure 1.**
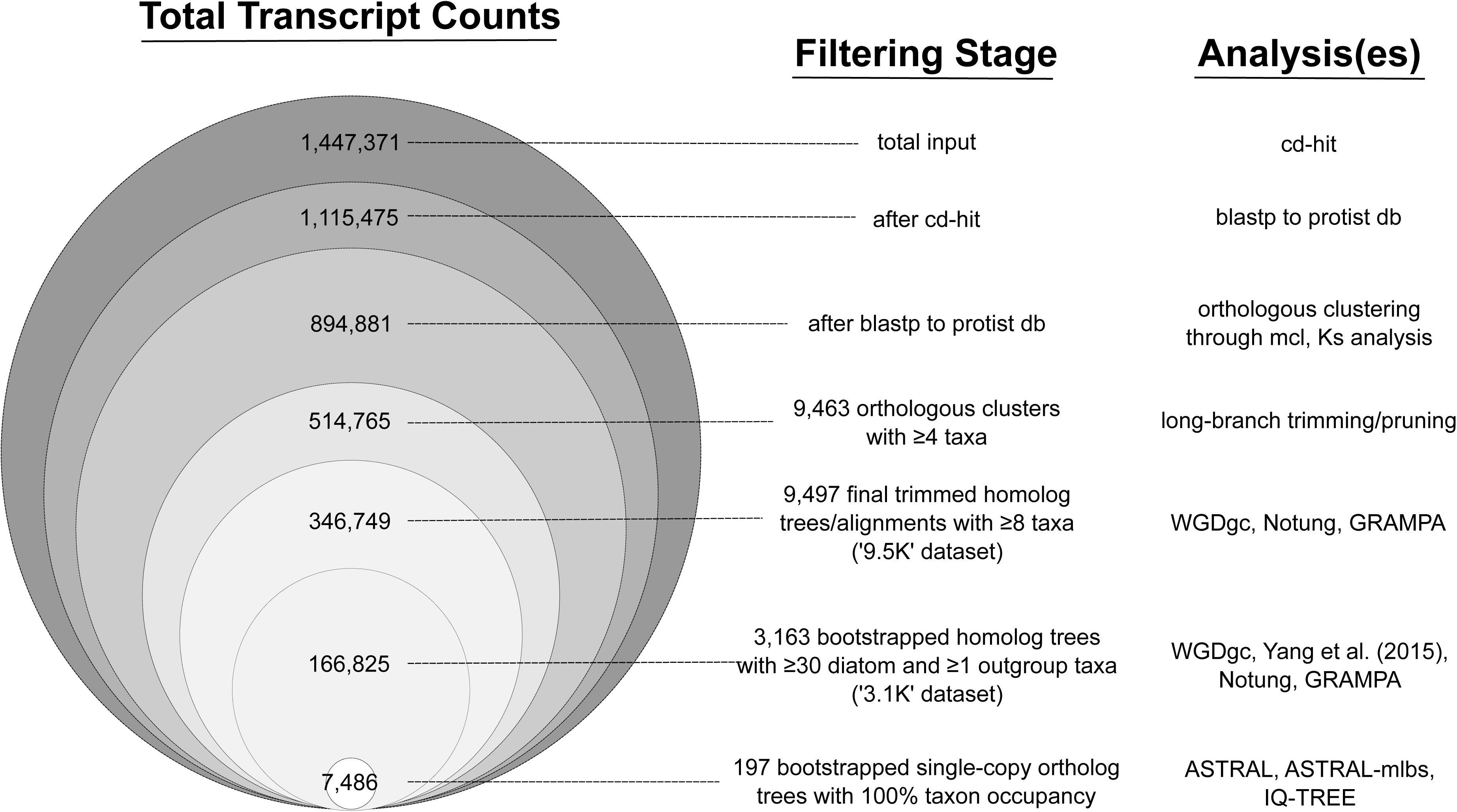
Dataset sizes at critical stages of analysis. The area of each circle is proportional to the total transcript count at that stage of analysis. Total transcript counts represent all assembled transcripts (transcriptomes) and predicted genes (genomes) available from all taxa at a given stage of analysis.

### Synonymous divergence (Ks) of paralogs

We looked for evidence for historic genome duplication events based on pairwise divergence between paralogs at synonymous sites (Ks) in both diatom and outgroup genomes (Lynch and Conery, 2000; Blanc and Wolfe, 2004). Methods for identifying secondary Ks peaks can vary considerably in several parameters (e.g., clustering criteria for paralogs and codon substitution model), and the behaviors of different Ks pipelines have not been systematically evaluated, so we used several different available pipelines and settings. We restricted Ks analyses to a set of relatively conserved genes, based on a BLASTP search (e-value ≤ 10^-10^) of each transcriptome against a database of complete proteomes from 17 protist species. The first approach followed Johnson et al. (2016), with initial filtering of each gene set to remove highly similar sequences (e.g., isoforms or very recent duplicates) using CD-HIT-EST (-c 0.98 -aS 0.90). Remaining proteins were then clustered for each species with CD-HIT (-c 0.40 -aL 0.75 -n 2), aligned with MAFFT, and back-translated by forcing nucleotide sequences to protein alignments with Pal2Nal (Suyama, Torrents, and Bork, 2006), with gap regions and internal stop codons removed. For each pair of paralogous nucleotide sequences in the CD-HIT clusters, Ks was calculated using the KaKs_Calculator (Zhang et al. 2006) under both the YN (Yang and Nielsen, 1998) and GY (Goldman and Yang, 1994) codon substitution models. We also estimated Ks distributions using the FASTKs pipeline (McKain et al., 2016) with default settings. The Trinity transcriptome assembler distinguishes closely related paralogous genes from isoforms of the same gene (Grabherr et al., 2011a). As a result, transcript assemblies are hierarchically organized according to assembly read clusters, which are comprised of ‘gene’ and gene ‘isoforms’. In some cases, isoforms of the same Trinity ‘gene’ might represent recently diverged paralogs, and some Ks pipelines are ‘Trinity-agnostic’, instead relying on alternative filtering strategies to distinguish paralogs and isoforms (Jiao et al., 2011; Johnson et al., 2016). Due to this ambiguity, Ks distributions were determined using the FastKs pipeline both before and after removing BLASTN self-hits at the ‘gene’ level for the Trinity assemblies (i.e., BLAST hits between two Trinity isoforms of the same Trinity gene). For both pipelines, we tested for multiple normal distributions in the Ks distributions using the R package mclust (Fraley et al., 2012), with the best fit model chosen using the Bayesian Information Criterion (BIC).

### Gene-tree reconciliation

#### Focal points of gene duplication and loss

We applied two gene-tree reconciliation strategies to two subsets of homolog trees to identify parts of the species tree with concentrations of gene duplication (and loss) events. First, we applied the approach used by Yang et al. (2015) to a set of 3163 homolog alignments (‘3.1K dataset’) filtered to include at least 30 diatoms and one outgroup taxon (Fig. 1). This pipeline maps rooted clades of homolog (orthologs and paralogs) trees to a species tree to determine the proportion of duplicated gene families, taking into account confidence in homolog tree topologies as measured by average bootstrap support across a sampled clade. For this analysis, we used RAxML, with the PROTCATWAG model and 100 bootstrap pseudoreplicates, to reconstruct homolog trees. Average bootstrap support values were relatively low across homolog trees, so the bootstrap cutoff was set at 40% (Yang et al., 2015).

Second, we used Notung (ver. 2.9) (Durand, Halldorsson, and Vernot, 2006; Darby et al., 2017) to reconcile and root two sets of gene trees: the set of 3163 homolog trees (‘3.1K dataset’, as described above) and a broader set of 9497 homolog trees with at least 8 diatoms (‘9.5K dataset’) (Fig. 1). We ran Notung’s phylogenomic pipeline to estimate the number of gains and losses in each gene tree and total counts of duplication and loss per node for the entire set of 9497 homolog trees. For the 3.1K dataset, we also performed bootstrap-based rearrangements, which minimize the reconciliation cost by making rearrangements around poorly supported nodes. We applied three bootstrap thresholds (40%, 50%, and 70%) and repeated the Notung phylogenomic pipeline on each of the resulting sets of rearranged trees. We applied relatively low bootstrap thresholds because the overall levels of bootstrap support in gene trees were low, i.e., only about 30% of nodes across all trees had bootstrap values >50%. In addition to bootstrap-based rearrangement, we also run Notung with gene trees that had average bootstrap support > 50% (total=374).

#### WGD validation at duplication focal points

To specifically test for the mechanism of WGD formation at focal nodes highlighted by the Yang and Notung pipelines, we used the software package GRAMPA (Thomas, Ather, and Hahn, 2017) to compare the reconciliation scores of multiply-labeled (MUL) trees against the singly-labeled species tree using homolog trees from the 3.1K and 9.5K datasets. Cases when the MUL tree – a topology in which a taxon or clade appears twice as the result of a duplication – had a better reconciliation score than the species tree were considered supportive of a WGD event. By default, GRAMPA performs least-common ancestor (LCA) reconciliation of all gene trees against both the species tree and all possible MUL trees, and reports the number of duplications and losses and their sum (the reconciliation score). Overly complex gene trees, which might take a prohibitively long time to reconcile, are filtered out based on a maximum allowed number of polyploid groups, which we set to 12 (GRAMPA’s group cap setting, default=8).

We ran GRAMPA with two basic strategies. First, we made no assumptions about the placement of polyploid lineages by excluding GRAMPA’s H1 and H2 parameters. We refer to these analyses as “unconstrained”. This approach tested all possible arrangements for the two parents of a putative allopolyploid event, including the same parent for autopolyploid events. For these analyses, a substantial fraction of gene trees were also filtered out due to the group cap setting (36% for the 3.1K dataset without rearrangements and 20% for the 9.5K dataset). To minimize this filtering and to base our inferences on the largest possible sets of trees, we also ran GRAMPA for each tested internal and terminal node separately by setting the H1 node and letting GRAMPA find the best H2 node or nodes. This reduced the number of alternative MUL topologies to those relevant for the focal node and resulted in the filtering out of many fewer trees as overly complex given our group cap setting of 12 (maximum of 15% and 16% for the un-rearranged 3.1K and 9.5K datasets, respectively). These analyses are subsequently referred to as “constrained”. We ran these analyses for all datasets, including the unrearranged 3.1K and 9.5K sets of trees, the rearranged versions of 3.1K trees at three bootstrap thresholds (40%, 50%, and 70%), and the pre-filtered set of trees with mean bootstrap >50%. Although for each dataset the “constrained” and “unconstrained” tests started with the same set of gene trees, depending on the topology of the relevant MUL trees, GRAMPA filtered out different sets of gene trees as overly complex. The reconciliation scores between the two search strategies and the scores of runs with different focal (H1) nodes are therefore based on slightly differing sets of input trees and are not comparable.

### Gene count analyses

We further tested 18 inferred duplication events identified from Ks distributions and gene-tree reconciliation methods (located on 11 terminal and 7 internal branches) with gene-count data derived from both the 3.1K and 9.5K datasets (Appendix 4) using the R package WGDgc (Rabier, Ta, and Ane, 2014). Initial tests used the entire species phylogeny, and required an orthologous cluster to include *Triparma pacifica* and at least one ingroup species, thereby removing orthologous clusters unique to diatoms. Using this strategy, most of the putative WGD events identified through Ks analyses were not detectable, likely due to excessively stringent filtering to meet the above criterion. Similar results have been observed in other studies that use gene count data, and one common solution is to focus analyses on subtrees that maximize the amount of data relevant to testing a particular WGD hypothesis (Tiley, Ane, and Burleigh, 2016). To increase the pool of orthologous clusters for detection of WGD events, while keeping computation memory and time reasonable, we created datasets and pruned accordingly the time-calibrated chronogram to include only those taxa relevant to a specific WGD hypothesis. For example, when testing the putative Ks-inferred WGD in *Gyrosigma*, we pruned the species tree down to include raphid pennates only (Fig. 2). The final datasets represented orthologous clusters represented in the outgroup and at least in one species of the ingroup. WGDgc analyses were run with the root prior set to the mean number of copies per cluster in each of the datasets and with the option “oneInBothClades” that reflected our filtering strategy. The putative WGD events were assumed to have occurred at the midpoint of branches leading to the focal node. Hypotheses were tested using likelihood ratio tests against a null model of no WGD events (Rabier, Ta, and Ane, 2014; Tiley, Ane, and Burleigh, 2016).

**Figure 2.**
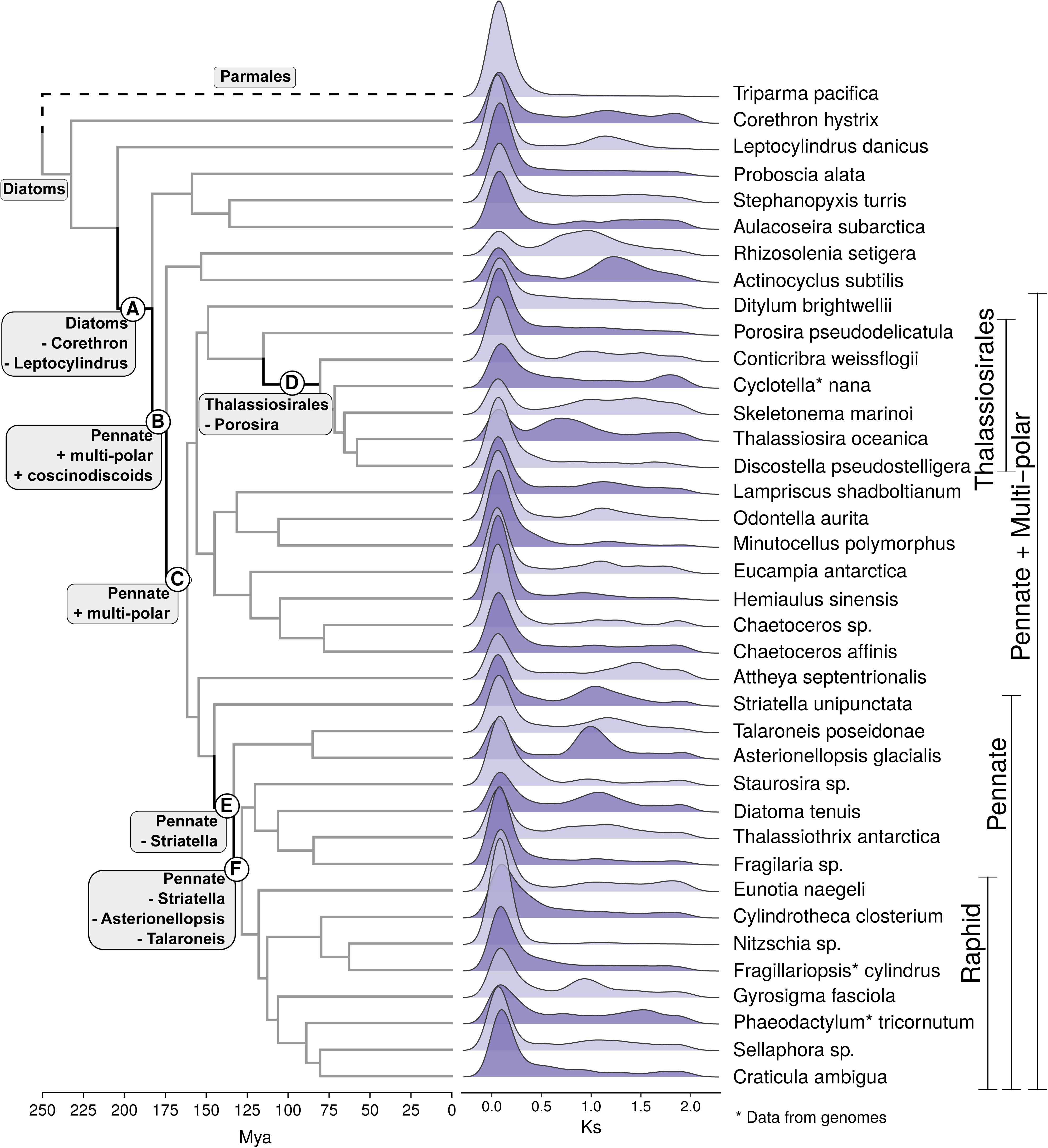
Time-calibrated species tree of 37 diatoms and the outgroup *Triparma* (Parmales) reconstructed from a concatenated alignment of 197 single-copy genes. Nodes relevant to downstream analyses are labeled (A–F). Ks-based age distributions were calculated with CD-HIT filtering and the GY model of codon substitution.

## RESULTS

### Assembly results

A total of 34 diatom and one outgroup (*Triparma pacifica*) taxa were assembled from paired-end RNA-seq read pools ranging in size from 21.3 to 424 million reads. Trinity assemblies ranged in size from 13 578 to 61 091 genes and 16 145 to 70 488 transcripts (including isoforms). BUSCO recovery averaged 70 ± 8% for combined complete and fragmented orthologs. Gene counts for protein sets from the five genome sequences ranged from 10 402 to 27 137 genes, with a corresponding average BUSCO recovery of 83 ± 6%. Sample information and assembly details are available in Appendix 1.

### Homology and orthology inference

A total of 9463 orthologous clusters containing at least four taxa were circumscribed with MCL (Fig. 1). After branch-length-based pruning, 9497 alignments and corresponding phylogenetic trees with at least eight taxa were recovered. These alignments were then filtered based on various taxon-occupancy thresholds to create data subsets for further analyses (Fig. 1).

### Species tree reconstruction

197 single-copy ortholog alignments with 100% taxon occupancy were recovered (Fig. 1), representing a combined alignment length of 58 294 amino acids. Coalescent-summary (ASTRAL, ASTRAL-mlbs) and concatenation-based (IQ-TREE) inference methods recovered generally well-supported species trees with identical branching orders, with the exception of the polar centric diatom *Ditylum brightwellii* (Fig. 2, Appendix 2), which was also difficult to place in another phylogenomic dataset (Parks, Wickett, and Alverson, 2017). Similar to previous findings (Parks, Wickett, and Alverson, 2017), relationships among the major multi-polar centric clades were the least supported in ASTRAL and ASTRAL-mlbs analyses. Gene-tree support varied across the species tree and, as in previous phylogenomic analyses of diatoms (Parks, Wickett, and Alverson, 2017), relationships among the major multi-polar clades were the most difficult to resolve, with deep splits supported by few or no gene trees (Appendix 2).

### Synonymous divergence (Ks) between paralogs

Ks-based age distributions of gene duplicates revealed secondary Ks peaks consistent with historic duplication events in most diatom species (Fig. 2), though the strength of the signal and the locations (average synonymous divergence) of secondary peaks varied by method, codon substitution model, and whether blastn self-hits at the Trinity ‘gene’ level were included in the analysis. The Ks distributions inferred from CD-HIT clusters (Johnson et al., 2016) using two substitution models (GY and YN) were largely overlapping, although the model used caused the size or placement of secondary peaks to shift slightly (to higher Ks values for GY model). Secondary Ks peaks inferred from BLAST-based clusters (McKain et al., 2016) were more distinct when blastn self-hits at the Trinity ‘gene’ level were removed and also tended to be both smaller and centered on lower Ks values than those called from CD-HIT clustering (Appendix 3). Two sister groups, *Actinocyclus subtilis* + *Rhizosolenia setigera* and *Asterionellopsis glacialis* + *Talaroneis poseidonae*, with strong secondary Ks peaks were each sister taxa on the species tree, possibly indicative of shared duplication events in those clades (Fig. 2). Secondary peaks in two other pennate diatoms, *Striatella* and *Diatoma*, suggest either a deeper, shared WGD along the pennate backbone or independent WGDs in these taxa (Fig. 2); the secondary Ks peaks in these two taxa were not, however, recovered by all of the Ks-based methods. Although less striking than those highlighted here, mclust identified numerous secondary Ks peaks in several other taxa as well (Appendix 3).

### Gene-tree reconciliation (Yang and Notung pipelines)

Although gene-tree reconciliation results largely agreed across analyses and sets of gene trees, bootstrap-based filtering and gene-tree rearrangement had a substantial impact on the number of families with inferred duplications and losses (Fig. 3). The Yang and Notung gene-duplication pipelines highlighted six branches along the backbone of the species tree with high concentrations of gene duplications (Figs. 2, nodes A-F). Four of these branches retained a high percentage of gene duplications irrespective of the set of gene trees used: the 9.5K and 3.1K sets for Notung and the 3.1K set for the Yang pipeline (Fig. 3). These nodes were: (1) the MRCA of all diatoms excluding *Corethron hystrix* and *Leptocylindrus danicus* (‘branch A’), (2) the MRCA of pennate and multipolar centric diatoms (‘branch C’), (3) the MRCA of Thalassiosirales excluding *Porosira pseudodenticulata* (‘branch D’), and (4) the MRCA of all pennate diatoms excluding *Striatella unipunctata* (‘branch E’) (Fig. 3). Aside from the three deepest nodes on our species tree (MRCA of *Triparma* + diatoms, the MRCA of diatoms, and the MRCA of all diatoms except *C. hystrix*), all other nodes across the species tree featured moderate to high proportions of gene loss, including the six nodes identified with high rates of duplication (Fig. 3).

**Figure 3.**
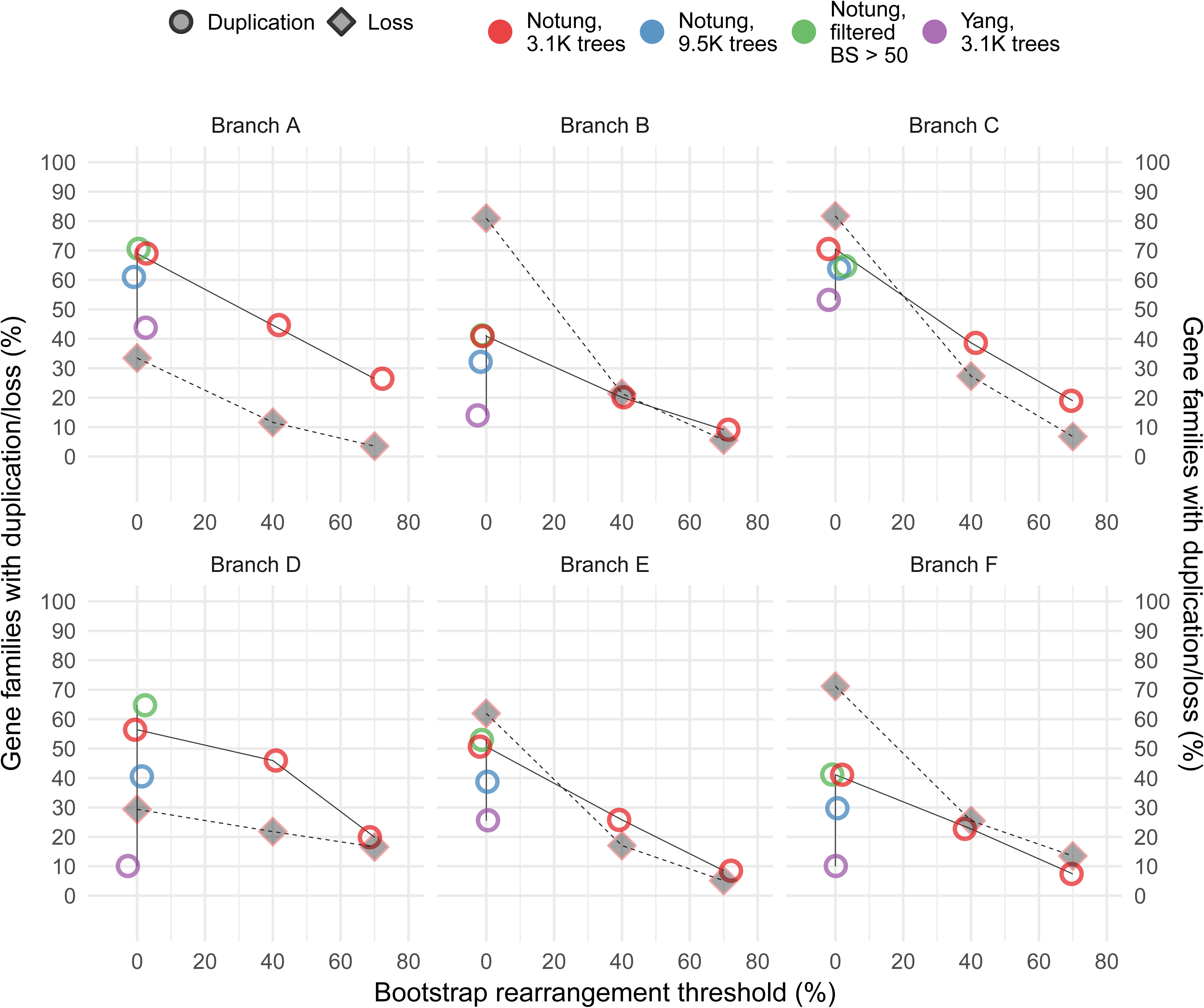
Trends in gene duplication and loss across select nodes of species phylogeny. Percent gene duplication and loss at six nodes discussed in the text were reconstructed with different sets of homolog trees, different reconciliation pipelines, and at different bootstrap thresholds for rearrangement. Refer to Figure 2 for a key to the branch names.

We also used Notung to perform bootstrap-based rearrangements of gene trees in order to conservatively estimate counts of gene duplications and losses. For these rearrangements, nodes in gene trees with bootstrap support lower than a set threshold that were inconsistent with the species tree were rearranged to minimize the number of inferred duplications and losses. Analyses of rearranged trees reduced the number of families with duplications and losses across all internal nodes, including the six focal nodes identified by the Yang and Notung pipelines and shifted a small portion of the events towards the tips of the tree. The number of gene families with inferred duplications and losses also decreased, as expected, as the bootstrap threshold was increased (Notung analyses with 3.1K dataset; Fig. 3). Among the six Yang/Notung focal nodes, reconciliation of the original gene trees found duplications in some 40–70% of gene families and losses in as many as 80% of gene families (Fig. 3). With a bootstrap threshold of 40%, the maximum percent of duplicated families reduced to ∼50% and further decreased to ∼26% at a bootstrap threshold of 70% (Fig. 3). The percentage of gene families with inferred losses dropped even more precipitously between the sets of original and rearranged trees, reducing from ∼80% to 20–30% for the most loss-rich nodes when rearranging at a 40% bootstrap threshold (Fig. 3). Duplication and loss counts continued to drop as the bootstrap threshold was increased further (Fig. 3). Overall, even at the most stringent bootstrap rearrangement threshold, the top three nodes had ≥19% of gene families with duplications, suggesting that gene tree reconciliation detected some signal for synchronous duplication events at or near these nodes (branches A, C, D; Fig. 3).

### Gene-tree reconciliation (GRAMPA)

These analyses were designed to test more specifically whether the high concentrations of duplicated gene families at focal branches identified by the Yang/Notung pipelines were due to WGDs. Our GRAMPA searches detected pervasive WGD signal in the absence of bootstrap-based rearrangement of the gene trees. For example, the least conservative, un-rearranged set of trees (9.5K dataset) recovered 481 MUL trees in unconstrained analyses, involving 42 different polyploid clades (H1 nodes) that were better than the species tree by at least 1000 units. For the set of un-rearranged trees from the 3.1K dataset, there were 179 MUL trees in unconstrained analyses, involving 27 different polyploid clades that scored better than the species tree by at least 1000 units. Nearly all detected signal was, however, very sensitive to gene tree support, with just two WGD events supported with a bootstrap rearrangement threshold of 40% [constrained analyses of branches A and C (Fig. 2)]. Nonetheless, resampling based on bootstrap-filtered gene trees, and clade-based resampling of both gene tree subtrees and MUL trees, revealed considerable support for WGD at several focal branches, including within the Thalassiosirales clade (Fig. 2, branch D). These results are described in detail in the following sections.

#### Whole-genome duplication at deep internodes

The Yang/Notung reconciliations against the singly labeled species tree inferred high concentrations of duplicated families at the MRCA of all diatoms excluding *Leptocylindrus* and *Corethron* (Figs. 2 and 3, branch A), and at the MRCA of pennate+multi-polar diatoms (Figs. 2 and 3, branch C). The former clade, all diatoms except *Leptocylindrus* and *Corethron* (Fig. 4), was not among the highest supported in GRAMPA analyses, although this scenario was still better than assuming no polyploidy. With unrearranged gene trees, GRAMPA detected the strongest signal for a putative WGD involving the latter clade (branch C, Fig. 4), and a GRAMPA search constrained to this clade only, found similar results using gene trees rearranged at a 40% bootstrap threshold.

**Figure 4.**
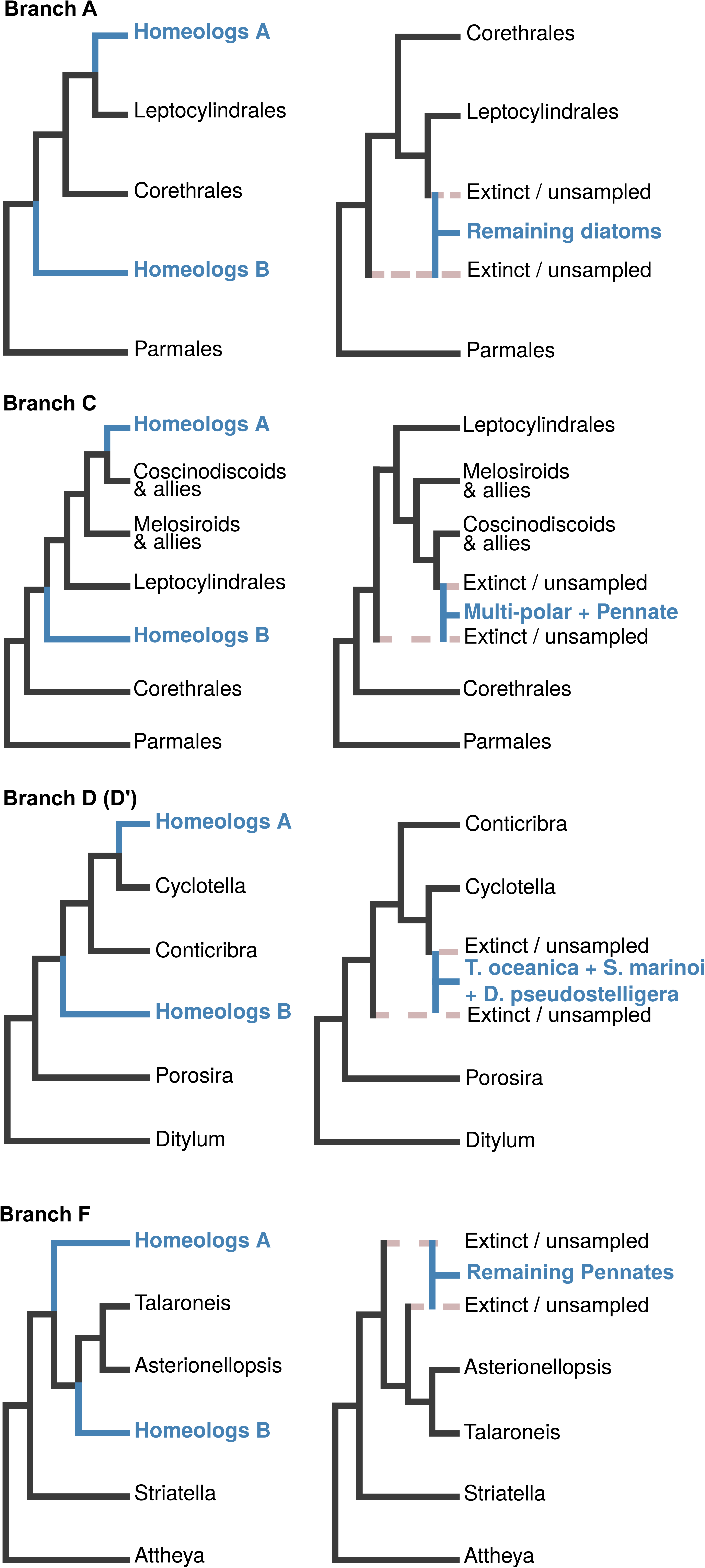
Results of GRAMPA reconciliation against multiply labeled (MUL) trees. Trees on the left represent the best MUL trees for each focal clade, with the corresponding network representation of the inferred allopolyploid events shown on the right. For the best MUL trees (left), the two sets of homeologs are indicated in **Blue**. In each case, the species tree placement of the allopolyploid clade is shown as ‘Homeologs A’; the placement of the second subgenome of the allopolyploid is marked as ‘Homeologs B’. For inferred allopolyploid events (right), parental and allopolyploid lineages are indicated in **Grey dashed** and **Blue solid** lines, respectively. In all cases, the inferred parental taxa of the allopolyploid clades are either extinct or have not been sampled. Branch names correspond to labeling in Figure 2.

In unconstrained GRAMPA searches, the best-ranked MUL tree involving all diatoms except *Leptocylindrus* and *Corethron* was 3919–14 893 units worse than the overall best MUL tree (MRCA of pennate+multi-polar diatoms), and MUL trees involving this clade were never better than the species tree when gene trees were rearranged. However, GRAMPA runs constrained to MUL trees specific to the MRCA of all diatoms excluding *Leptocylindrus* and *Corethron* found support for a WGD event up to 40% bootstrap rearrangement threshold, with reconciliation scores at least 407 units better than the species tree. This outcome was likely due to more gene trees passing GRAMPA’s tree-complexity filter when the search was constrained to fewer MUL trees, in this case only those relevant for the clade of all diatoms excluding *Leptocylindrus* and *Corethron*.

We repeated the Notung and GRAMPA analyses focusing on the above clades and using only trees with average bootstrap support ≥50% (total=374) filtered from the 3.1K dataset (Fig. 3). In this case, the best MUL tree was 415 units better than the singly labeled species tree, and again identified an allopolyploidy event involving the clade of pennate + multi-polar diatoms and an extinct or unsampled second parental lineage represented with the ancestor of all diatoms except *Corethron* (branch C, Fig. 4). MUL trees involving the MRCA of pennate + multi-polar + coscinodiscoid diatoms, multi-polar diatoms (minus *Attheya*), and the MRCA of all diatoms except *Leptocylindrus* and *Corethron* also scored high. Overall, results based on this set of well-supported gene trees agreed with the results from our analyses of the entire sets of un-rearranged gene trees, suggesting that rearrangements, even at low bootstrap thresholds, might be overly conservative. The Yang et al. (2015) pipeline, which extracts and calculates duplication counts on clades with average bootstrap beyond a certain threshold, gave similar results.

#### Allopolyploidy within Thalassiosirales

In unconstrained searches, GRAMPA found that MUL trees defined by the branch representing Thalassiosirales – *Porosira pseudodenticulata* (Figs. 2 and 4, branch D) were never better than the species tree. This applied irrespective of the set of trees used and whether or not the trees were rearranged. This result was also confirmed by an analysis of only the Thalassiosirales subtree with *Ditylum brightwellii* as the outgroup. Closer examination of other MUL trees relevant to Thalassiosirales revealed several smaller clades nested within the Thalassiosirales – *Porosira* clade (Fig. 5, branch D) with strong support for a WGD (branch D/D′, Fig. 4). The strongest support was observed for a clade comprised of *Thalassiosira oceanica*, *Skeletonema marinoi* and *Discostella pseudostelligera* (branch D′, Figs. 4 and 5), with their MRCA as one parent and the MRCA of all Thalassiosirales excluding *Porosira* as the second parent of an allopolyploid event. These results were robust to the different sets of trees used, analyses performed at different phylogenetic scales (all diatoms or Thalassiosirales only), and to bootstrap-based rearrangement up to a threshold of 50%. To further evaluate these results, we extracted the Thalassiosirales + *Ditylum* clade from all trees in the 3.1K dataset and filtered the resulting subtrees to: (1) include a minimum of three species, and (2) have an average bootstrap support ≥70. A total of 240 gene trees met these criteria. Repeating the reconciliation analyses with Notung and GRAMPA on this conservative set of well-supported trees returned identical results, supporting the clade comprised of *T. oceanica*, *S. marinoi* and *D. pseudostelligera* as allopolyploid (branch D/D′, Fig. 4). Such outcomes, i.e., a concentration of gene duplications in a branch older than the MRCA of the polyploid clade, have been interpreted as strong support for ancient hybridization and allopolyploidy in yeast (Marcet-Houben and Gabaldon, 2015; Thomas, Ather, and Hahn, 2017). The discrepancy between the relative age of the reconciliation-inferred peak of duplications (Fig. 5, branch D) and the polyploid clade (Fig. 5, branch D′) is therefore likely due to the earlier divergence of the hybridization-derived homeologs present in the genome of the polyploid lineage, which trace back to the earlier branch D, compared to the age of the polyploid species lineage itself.

**Figure 5.**
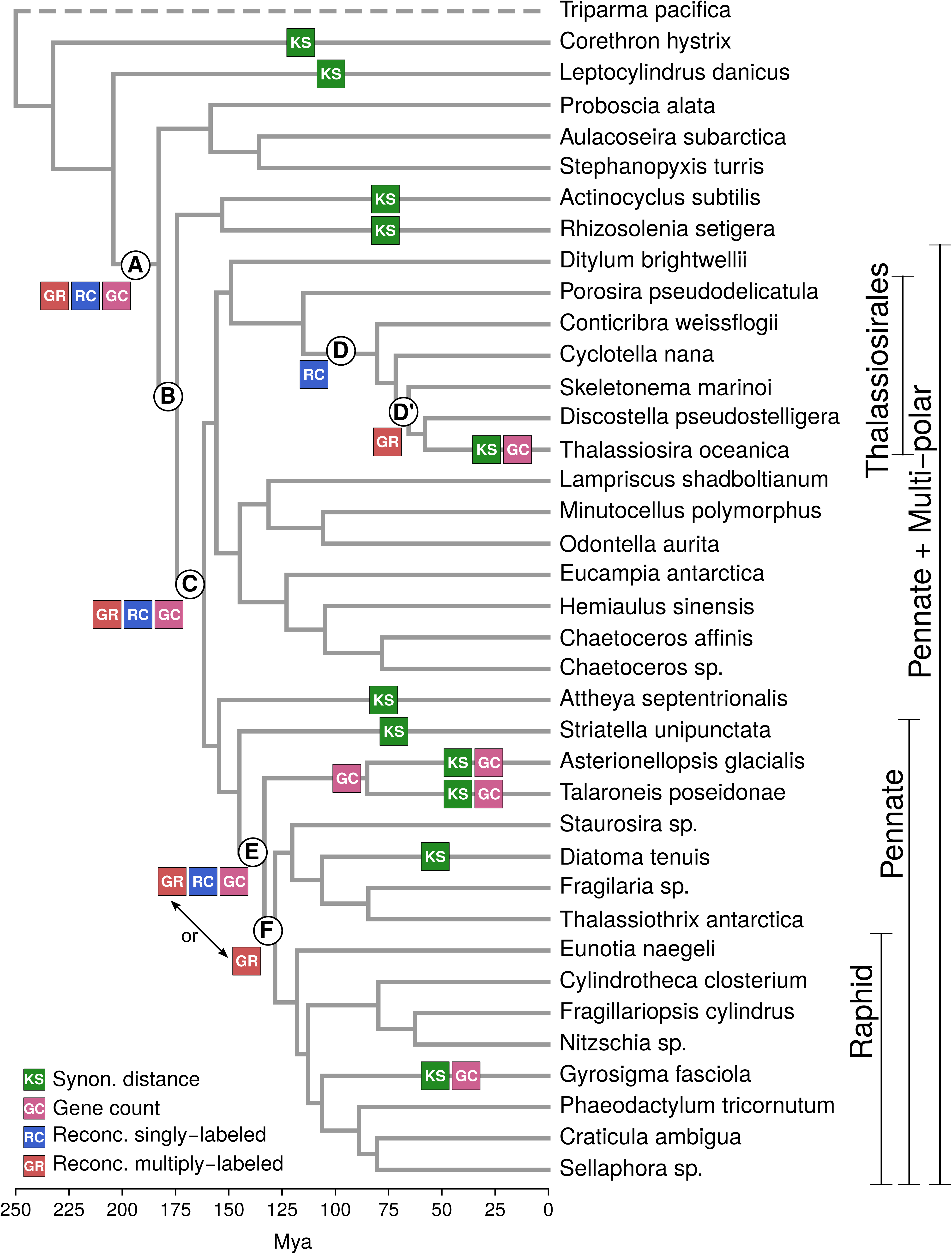
Summary of WGD across diatoms. **Green**: WGDs supported by Ks-based age distributions of duplicated genes (KS). **Pink**: WGDs detected with gene count data (GC). **Blue**: WGDs inferred by reconciliation of gene trees against the singly labeled species tree (RC). **Red**: WGDs inferred by reconciliation of gene trees against multiply labeled trees (GR). Branches discussed in the text are labeled (A–F). Within Thalassiosirales, D’ denotes a GRAMPA-inferred allopolyploid clade (GR) that did not coincide with the duplication peak inferred from the Notung and Yang analyses (D). Within pennate diatoms, GRAMPA-inferred events are added to both branches E and F to reflect uncertainty in the placement of the WGD.

#### Allopolyploidy within the pennate clade

The unconstrained GRAMPA searches found strong support for polyploidy of either the MRCA of pennate diatoms excluding *Striatella* (Fig. 2, branch E) or the clade of all pennate diatoms excluding *Striatella*, *Asterionellopsis,* and *Talaroneis* (Fig. 2, branch F). To investigate potential pennate-specific WGD events, we ran GRAMPA on the subtree containing only pennate diatoms, with *Attheya* as the outgroup. The best reconstructions identified the clade of pennates excluding *Striatella*, *Asterionellopsis*, and *Talaroneis* as the most likely polyploid clade (Figs. 2 and 4, branch F), with the second parental lineage being extinct or unsampled member of the clade subtended by the MRCA of all pennate diatoms excluding *Striatella*. Events specific to: (1) all pennate diatoms except *Striatella*, (2) the raphid pennate clade, and (3) a smaller clade comprised of *Diatoma*, *Fragilaria*, and *Thalassiothrix* had lower, but comparable scores, all of which were better than the species tree.

These results were not, however, robust to bootstrap rearrangements, with support for allopolyploidy disappearing after gene-tree rearrangement with a 40% bootstrap cutoff. Searches constrained to MUL trees defined by the MRCA of all pennates except *Striatella* (branch E, Figs. 2 and 5), or the subsequent clade [all pennates excluding *Striatella*, *Asterionellopsis*, and *Talaroneis* (branch F, Figs. 2 and 5)] showed similar results, with only the former clade robust to bootstrap rearrangement. However, an analysis of pennate diatom subtrees with a mean bootstrap support ≥70% (84 in total) again showed support for a polyploid clade composed of all pennate diatoms except *Striatella*, *Asterionellopsis*, and *Talaroneis*. Further, this analysis also supported a smaller clade within raphid pennates composed of *Sellaphora* and *Craticula* as polyploid. The best MUL tree in this case placed the second parental lineage of this inferred allopolyploid event at the MRCA of *Asterionellopsis* and *Talaroneis*, suggesting that the actual second parent might be an extinct or unsampled lineage from the clade circumscribed by the MRCA of *Asterionellopsis* + *Talaroneis* and all of its descendants (branch F, Fig. 4). Overall, although bootstrap-based rearrangement erased most of the signal for polyploidy within the pennate clade, analyses of a small set of strongly supported trees largely agreed with the inferences made from the entire set of un-rearranged phylogenies.

### Gene count analyses

Analyses based on gene counts were designed to test 18 putative WGD events whose placements were based either on analyses of paralog divergence at synonymous sites (10 terminal and 2 internal branches; Fig. 2) or gene-tree reconciliation (6 internal branches) (Figs. 4 and 5). Each putative event was tested independently through comparison to a null, non-WGD model. We performed the analyses using gene counts based on both the 3.1K and 9.5K datasets and, with three exceptions as described below, recovered the same set of results for both analyses. We detected WGDs in eight out of 18 tested branches, with relatively low rates of homolog retention following the duplication event, whereby the retention rate (*q*) is defined as the probability of retaining the WGD-derived copy of a gene (Rabier, Ta, and Ane, 2014). Retention of two WGD-derived homologs following a duplication event was generally <2%, though four WGDs had retention rates between 3% and nearly 15%. All eight tests that returned support for WGD via retention rates > 0 were significantly better than their corresponding no-WGD null models (likelihood ratio tests, df=1, *X*^2^ P-value ≤ 0.001for all tests; Appendix 4).

Within pennate diatoms, gene-count analyses detected the Ks-inferred WGDs in *Gyrosigma* (*q* = 14.8%), *Asterionellopsis* (*q* = 7.0%), and *Talaroneis* (*q* = 1.4%) (Fig. 5). There was also signal for WGD along the branch leading to the MRCA of *Asterionellopsis* and *Talaroneis* (*q* = 3.6%), suggesting that the Ks peaks observed in these two taxa might represent a shared WGD (Fig. 2, 4). Finally, in agreement with gene-tree reconciliation results, we also detected signal for WGD along the branch leading to the MRCA of all pennate diatoms excluding *Striatella* (*q* = 1.1%) (Fig. 5). The latter event was not detected with gene counts derived using the 3.1K dataset, which instead detected signal for WGD in *Attheya* alone (*q* = 0.3%). WGD along the branch leading to *Attheya* was not observed in the analysis of counts calculated from the 9.5K dataset..

Several nodes across the centric and multi-polar centric diatoms were also tested using WGDgc. We tested two distinct hypotheses within Thalassiosirales: the Ks-inferred WGD in *Thalassiosira oceanica* and the GRAMPA-inferred WGD at the MRCA of *Thalassiosira oceanica*, *Skeletonema marinoi*, and *Discostella pseudostelligera*. We detected signal for an event within the *T. oceanica* lineage (*q* = 1.2%) but found no evidence for the older WGD (Fig. 5). Despite distinct Ks peaks, we did not detect WGD events in *Actinocyclus*, *Rhizosolenia*, or their MRCA, nor did we find support for the WGD events implied by secondary Ks peaks in *Corethron* and *Leptocylindrus*. Finally, we tested for two events supported by both reconciliation and GRAMPA results, at the MRCA of pennate + multi-polar diatoms (Figs. 2 and 4, branch C) and the MRCA of all diatoms excluding *Corethron* and *Leptocylindrus* (Figs. 2 and 4, branch A). The gene-count analysis detected WGDs on both branches, including a WGD with retention rate *q* = 1.7% along the branch leading to pennate + multi-polar diatoms using both datasets and an event on the branch leading to the MRCA of all diatoms excluding *Corethron* and *Leptocylindrus* using the 9.5K dataset (*q* = 6.4%) (Fig. 5).

## DISCUSSION

Substantial variation in genome size and chromosome number, a high rate of genome size evolution, and direct observations of polyploidization in cell cultures together suggest that diatom genomes might have undergone past WGD events (Mann, 1994, 1999a; Oliver et al., 2007). Our survey of 37 diatom genomes and transcriptomes provided strong support for this hypothesis, identifying as many as 16 separate historic WGDs across diatoms, seven of which were supported by multiple lines of evidence. Our exemplar-based taxon sampling precluded precise pinpointing of the timing of these events, with four strongly supported events assigned to terminal branches that represent ca. 60-100 million years of evolutionary history. Nevertheless, despite our sampling and general challenges of working with a group of non-model organisms, our analyses point to a relatively extensive history of WGD in diatoms.

### Mechanisms of polyploid formation in diatoms

Although auto- and allopolyploids are equally abundant in angiosperms (Barker et al., 2016), the mechanisms underlying polyploid formation are much more poorly known in diatoms. Our results suggest that allopolyploidy may be especially important in diatoms, though the patterns and rates of hybridization are very poorly known. High sequence divergence in homologous chromosome assemblies from a raphid pennate diatom, *Fistulifera solaris*, points to an allodiploid origin of that species (Tanaka et al., 2015). Given the time- and labor-intensive nature of experimental reproductive studies of diatoms (see Chepurnov et al., 2004; Mann et al., 2004; Chepurnov et al., 2008; Chepurnov et al., 2012), evidence supporting hybridization and introgression in diatoms is likely to come from genomic data (Mallet, 2005), emphasizing the need for more intensive studies focused on taxon-rich clades at lower phylogenetic scales. Candidates for such studies include *Ditylum brightwellii* (Koester et al., 2010), *Sellaphora* (Mann et al., 2004; Evans et al., 2008), *Seminavis* (Moeys et al., 2016), *Pseudo-nitzschia* (Casteleyn et al., 2009; Basu et al., 2017), and *Cocconeis* (Geitler, 1927; Geitler, 1973). Given the evidence for relatively frequent ancient hybridization uncovered by our analyses, including in the pennate diatoms, it will be important to determine the specificity of sex pheromone systems used by pennate diatoms for mate attraction (Sato et al., 2011; Gillard et al., 2013; Moeys et al., 2016). Finally, although our analyses highlighted allopolyploidy as a potentially important mode of WGD in diatoms, it is important to note that autopolyploidy may be shown to be equally, if not more, common with increased sampling. Indeed, autopolyploid formation has been directly observed *in vitro* for several different species of raphid pennate diatoms (Geitler, 1927; Mann and Stickle, 1991; Mann, 1994; Chepurnov and Roschin, 1995; Chepurnov et al., 2002).

A number of observed meiotic anomalies suggest that diatom polyploids could form in a variety of ways. First, Meiotic nonreduction, which is thought to be the predominant mode of polyploid formation in plants (Thompson and Lumaret, 1992; Ramsey and Schemske, 1998), likely occurs in diatoms as well. Although the rate of meiotic nonreduction in diatoms is unknown, Mann (1994) observed that failed cleavage in gametangia of the raphid pennate diatom, *Dickea ulvacea*, led to the formation of ‘double gametes’ that produced a dikaryotic, triploid-like zygote following fusion with a reduced gamete (Mann, 1994). Second, although polyspermy is thought to occur relatively rarely in plants (Ramsey and Schemske, 1998), the production of triploid and tetraploid zygotes from simultaneous gamete fusions has been observed in culture studies of several raphid pennate diatoms (Geitler, 1927; Mann and Stickle, 1991; Mann, 1994; Chepurnov and Roschin, 1995; Chepurnov et al., 2002), suggesting that this may be a principal pathway to polyploidization in diatoms. These studies have found mixed populations of co-occurring haploid, triploid, and tetraploid zygotes following one or two rounds of crossing in culture, suggesting that two-step, ‘triploid-bridge’ routes to stable polyploidy may be more common in diatoms than other groups (Ramsey and Schemske, 1998). These hypotheses further underscore the value of the experimental reproductive studies in diatoms that initially led to these discoveries. Extending these studies to include longer-term tracking of *in vitro* polyploids will help clarify the long-term viability and reproductive dynamics of vegetative haploids, triploids, and tetraploids, thereby distinguishing culturing anomalies from observations that hint at the the natural frequencies and mechanisms of polyploid formation in diatoms.

### Combined genomic evidence for whole-genome duplication in diatoms

We applied three different strategies to a large transcriptomic dataset to investigate support for historical duplication events across the diatom lineage: (1) traditional Ks-based age distributions of duplicated genes; (2) tree-based reconciliation methods to identify nodes on the species tree with concentrations of gene duplications or to construct specific tests for allopolyploidy, and; (3) gene-count methods that provide conservative, sequence- and gene-tree-agnostic inferences of WGD. Although each of these approaches suffers some drawbacks, we considered a putative WGD as strongly supported when two or more analyses with disparate approaches were in agreement (Fig. 5).

Although Ks-based age distributions are useful for initial exploration of duplication signal within a genome, several clear challenges with these types of approaches became evident in diatom analyses. First, there is no consensus strategy for discerning duplication peaks from Ks distributions. In some cases, peaks are identified essentially ‘by eye’ (Blanc and Wolfe, 2004; Fawcett, Maere, and Van de Peer, 2009; Tang et al., 2010; Cannon et al., 2015), which can easily turn into an exercise in ‘the reading of tea leaves’. Although several statistical approaches have been adopted to identify discrete shifts or peaks in Ks distributions (Schlueter et al., 2004; Cui et al., 2006; Vanneste et al., 2015), secondary peaks may not always correspond to large-scale duplication events (Johnson et al., 2016) and there are no concise methods to distinguish between peaks representing WGD, large-segmental duplications, and small-segmental duplications based solely on Ks distributions. The identification of ancient duplication events is also challenged by saturation at synonymous sites (Vanneste, Van de Peer, and Maere, 2013), and this problem should be more pronounced in lineages with higher substitution rates. As unicells with short generation times, diatoms have relatively high rates of nucleotide substitution compared to multicellular lineages (Bowler et al., 2008). As a result, Ks-based age distributions are more likely to saturate sooner, erasing the signature of ancient WGDs (Vanneste, Van de Peer, and Maere, 2013). On average, 45% of the paralog pairs in a given species had Ks values that were out of range (>2) for drawing Ks-based inferences of WGD. Further analyses may show that these trace back to some of the deeper duplication events identified by gene-tree and gene-count analyses (Fig. 5).

In contrast to Ks plots, gene-tree reconciliation methods allow more rigorous statistical determination and increased confidence in phylogenetic mapping of large-scale duplication events (Durand, Halldorsson, and Vernot, 2006; Jiao et al., 2011), even allowing for specific tests of auto-vs. allopolyploidy (Thomas, Ather, and Hahn, 2017). The power of these approaches is limited, however, by the quality of the gene trees. Our gene trees were poorly supported in most cases. For the bootstrapped gene trees of our 3.1K dataset, the overall distribution of bootstrap values across all nodes of these trees was relatively low, with median bootstrap support = 29 for all gene tree nodes combined; 68% and 80% of the nodes across gene trees had bootstrap support lower than 50% and 70%, respectively. Bootstrap-based rearrangement of gene trees to minimize the numbers of inferred duplications and losses is a common strategy for guarding against false inferences of WGD from poorly supported gene trees (Durand, Halldorsson, and Vernot, 2006; Inoue et al., 2015; Thomas, Ather, and Hahn, 2017). All of our gene trees, and a majority of nodes within our trees, had the potential to be rearranged. Deciding on a bootstrap threshold on which to base our inferences, therefore, depended on our confidence in the correct reconstruction of the gene trees. The inference of WGD events should ideally be based on strongly supported nodes that, when reconciled against the species tree, identify duplication events (Hahn, 2007). However, the amount of data necessary to obtain strong support for nodes depends on tree shape and the distribution of internal branch lengths (Alfaro, Zoller, and Lutzoni, 2003; Hahn, 2007; Philippe et al., 2011). Short internal branches potentially require substantial amounts of data to obtain strong bootstrap support, but the amount of data (i.e., alignment length) available is clearly limited for individual gene trees. Simulation studies have shown even correct nodes can receive low bootstrap values under a variety of conditions (Alfaro, Zoller, and Lutzoni, 2003). These considerations highlight the difficulties in determining an empirical cutoff for what should be considered an accurate bipartition and, by extension, a bootstrap threshold for gene-tree rearrangement for reconciliation analyses.

Finally, empirical and simulation studies have shown that gene-count matrices also provide robust identification of WGD events (Rabier, Ta, and Ane, 2014; Tiley, Ane, and Burleigh, 2016), although these methods are conservative and may fail to identify multiple events in the same lineage and duplication events followed by high rates of gene loss (Hahn, 2007; Tiley, Ane, and Burleigh, 2016). We did not recover strong signal in any of our analyses for multiple duplication events in a single lineage; however, our analyses suggest low rates of duplicate retention are common to diatoms, as the majority of our tested branches featured very low gene retention rates (q<0.02). Similarly, our Notung analyses also identified relatively high rates of gene loss in nodes across the species tree (Fig. 3). This likely impedes the effectiveness of WGDgc in identifying WGD events in our dataset, and suggests that high rates of molecular and genome evolution in diatoms may rapidly mask signal from historic duplications and lead to underestimation of the number of duplication events. On the other hand, fast rates of gene loss following duplication coupled with WGDgc’s ignorance of gene tree topology and potential asymmetric gene loss, increase the confidence in WGDs inferred by gene count data. Our exemplar approach may also bias WGD events estimated from gene count data toward older events, as duplicated genes are more likely to be lost along the longer terminal branches of our species phylogeny. In this regard, denser taxon sampling may reveal gene-count support for putative terminal duplication events supported by Ks analyses.

### Ancient paleopolyploidy in diatoms

We identified two nested WGDs occurring roughly 200 Mya (Fig. 5, branch A) and 170 Mya (Fig. 5, branch C) that support polyploid ancestry for the vast majority of diatom diversity. Importantly, these were among the most robust WGDs in our analyses, being detected by gene tree reconciliation against the species tree, reconciliation against multiply-labeled trees, and by gene count analysis (Fig. 5). Although both events enjoy strong support from multiple analyses, limitations of our exemplar taxon sampling and lack of genome size and karyotype data for most of diatoms, impede a complete understanding of the precise origins of these events. For example, the inferred deep WGDs at branches A and C were supported by both reconciliation (Fig. 3) and gene count data (Appendix 4) and reconciliation against multiply-labeled trees clearly supported an allopolyploid mode of origin for both events. In both cases, the second parental lineage of the allopolyploid event was an extinct or unsampled lineage vaguely identified as the MRCA of all diatoms or all diatoms excluding *Corethron*. Although it is possible (or perhaps likely) that these ancestors are extinct, the fact that only two branches separate the older of these events and the diatom stem lineage leaves open the possibility that our sampling is too coarse for a precise determination of the lineages involved in this allopolyploid event.

Gene-tree reconciliations also showed some evidence for a polyploidy event in the MRCA of all diatoms, but several limitations of our dataset caution against over-interpreting this finding. With just a few dozen species (Ichinomiya et al., 2016), the sister lineage to diatoms (Bolidophyceae: Parmales) is a phylogenetic depauperon (Donoghue and Sanderson, 2015), resulting in long stem branches separating them from diatoms. The large number of morphological and life history differences distinguishing these two clades makes it difficult to polarize the large number of changes that have accumulated since they split roughly 200–250 Mya (Nakov, Beaulieu, and Alverson, 2017) **–** a limitation that extends to the gene family data used in our analyses. This problem is further exacerbated by the available Parmales data, which are currently limited to a single transcriptome from the flagellated, unsilicified stage of the life cycle. We know, for example, that the dataset is missing genes involved in silicification (Kessenich et al., 2014). A more complete representation of the Parmales genome will help show whether, like angiosperms, all diatoms have shared polyploid ancestry.

### Historic allopolyploidy in Thalassiosirales

Thalassiosirales is among the most common and abundant diatom lineages in the plankton of both marine and freshwaters. It is also a long-established, genome-enabled model system for studies of diatom physiology, morphology, and ecology (Guillard and Ryther, 1962; Armbrust et al., 2004; Poulsen and Kroger, 2004; Alverson, Jansen, and Theriot, 2007). The discovery of ancient hybridization and allopolyploidy in this group further establishes them as an excellent system for understanding these and other evolutionary processes in diatoms.

The signal for polyploidy within Thalassiosirales was among the strongest recovered in our analyses, being detectable even after applying a relatively stringent (given our gene trees) bootstrap rearrangement threshold of 50%. Gene tree reconciliation supported an allopolyploid event involving the clade comprised of *T. oceanica*, *S. marinoi* and *D. pseudostelligera* (Fig. 5, branch D′). Uncertainty in the species tree, however, makes it difficult to accurately circumscribe this event. Although most nodes within the Thalassiosirales species tree were well supported, gene tree discordance was especially high for splits within the putatively polyploid subtree [(*S. marinoi*, (*T. oceanica*, *D. pseudostelligera*))] (Appendix 2). Interestingly, the two nodes immediately predating this clade (the MRCA of Thalassiosirales minus *Porosira* and the MRCA of Thalassiosirales) had many more concordant gene trees, suggesting that the high levels of discordance in (*S. marinoi*, (*T. oceanica*, *D. pseudostelligera*)) may reflect, at least in part, conflict resulting from past hybridization (Appendix 2). Densely sampled phylogenies of Thalassiosirales inferred from ribosomal RNA and chloroplast genes produce an alternative topology [((*T. oceanica*, *S. marinoi*), *D. pseudostelligera*)], with *T. oceanica* sister to *Skeletonema* (Alverson, Jansen, and Theriot, 2007). If this relationship is correct, the strong secondary Ks peak in *T. oceanica* and the heavily tailed Ks distribution in *S. marinoi* (Fig. 2) raise the possibility the polyploid lineage is comprised of these two lineages alone, with the inclusion of *Discostella* representing an artifact of sparse taxon sampling and uncertainty in the species tree.

Finally, secondary peaks are evident in the age distributions of both *T. oceanica* and *S. marinoi*, and although less pronounced, age distributions of *Cyclotella nana* and *Conticribra weissflogii* also have heavy right tails (Fig. 2, Appendix 3). Although WGD inferences from Ks distributions alone, especially in lineages with high substitution rates, can be problematic (Vanneste, Van de Peer, and Maere, 2013 and see above), these results may nevertheless point to additional paleopolyploid events in the evolutionary history of Thalassiosirales, a hypothesis that has some support from *T. oceanica* data. Here, gene tree reconciliation supported a shared, deep WGD event, gene count analysis detected a *T. oceanica*-specific WGD, and its Ks distribution was one of two with readily observable secondary peaks (Fig. 2; Appendix 3). We cannot, therefore, rule out the possibility that the *T. oceanica* genome carries signal from two temporally distinct WGD events – its genome representing the product of as many as four paleopolyploidy events (Fig. 5).

### Historic allopolyploidy in pennate diatoms

The transition from radial to axial cell wall symmetry and from oogamous to isogamous sexual reproduction are landmark events in diatom evolution (Round, Crawford, and Mann, 1990), circumscribing a clade whose species diversity vastly outnumbers the remaining diatoms (Guiry and Guiry, 2017) and, as a result, motivating great interest in identifying the underlying drivers of this disparity. The clade of actively motile, raphe-bearing species nested within pennate diatoms (Fig. 2) diversified at a faster rate compared to the grade of clades of non-motile diatoms with axial and radial symmetry (Nakov, Beaulieu, and Alverson, 2017), suggesting that the evolution of active motility allowed these species to better utilize complex benthic habitats, leading to finer niche partitioning and faster rates of evolution and diversification (Nakov, Beaulieu, and Alverson, 2017). However, other factors have almost certainly influenced the diversification of pennate diatoms as a whole, and the results presented here point to a potential role for whole genome duplication.

Taking our combined results from gene tree reconciliation, gene count, and Ks analyses, we found evidence for as many as six independent WGD events within pennate diatoms (Fig. 5). Three of these events were supported by at least two out of the three strategies, whereas the others were more tenuous and observed only in the Ks-based age distributions (Fig. 5). The best-supported events included: (1) a deep split within the pennates circumscribing nearly the entirety of the clade (Fig. 5, branch E or F); (2) a deep split within the deepest clade of araphid pennates (Fig. 5, the MRCA of *Asterionellopsis* and *Talaroneis*), and; (3) a terminal branch representing the highly diverse navicuoloid diatoms with a stem age of > 100 Mya (Fig. 5, *Gyrosigma*). Placements of these events suggests that the majority of pennate diatoms share an ancient WGD, followed by multiple rounds of additional, nested polyploidizations that have affected several subclades of pennate diatoms (Fig. 5). Note also that these pennate-specific events might have occurred in addition to at least two earlier WGDs (Fig. 3, branches A and C), analogous to the complex polyploid ancestry of numerous angiosperm lineages (Bowers et al., 2003; Jiao et al., 2011).

As with WGDs in other parts of the phylogeny, a degree of uncertainty exists with regard to the deep, nearly pennate-wide WGDs (Fig. 5, branch E or F). Specifically, reconciliation against the species tree and gene count analyses indicated that the most likely placement of this event was at the branch representative of the MRCA of all pennate diatoms excluding *Striatella* (Fig. 5, branch E). Reconciliation against MUL trees was more equivocal, however, supporting either this branch or the next branch up the backbone as the likely ancestor that experienced the duplication (Fig. 5, branch F). As before, we were unable to determine whether this uncertainty is a byproduct of our exemplar sampling, i.e., the lineages relevant for pinpointing the placement of this event might be missing from our dataset. Alternatively, uncertainty in the species tree might be carried over into the placement of this WGD. More densely sampled phylogenies based on conventional phylogenetic markers place *Striatella* (along with *Asterionellopsis* and *Talaroneis*) within one of the clades in the araphid grade (Theriot et al., 2015; Nakov, Beaulieu, and Alverson, 2017). Phylogenomic analyses with fewer species but more markers place *Striatella* as sister to all other pennate diatoms (Fig. 2 and Appendix 2; see also Parks, Wickett, and Alverson, 2017). These competing hypotheses have clear implications for our ability to infer the location and timing of this WGD and further highlight this part of the tree as a primary target for additional genomic sampling.

Similar uncertainty, linked again to the granularity of our sampling, is evident in the (*Asterionellopsis* + *Talaroneis*) clade. Ks-based age distributions highlighted strong secondary peaks in these two taxa (Fig. 2, Appendix 3), possibly indicative of a shared WGD in the MRCA of this clade. We found evidence to support this hypothesis as well as signal for WGD on each of the terminal branches leading to *Asterionellopsis* and *Talaroneis* (Fig. 5, Appendix 4). A parsimonious interpretation would support that all of this signal traces back to the shared event, but given the stem ages of these lineages (∼80 Mya; Figs 2 and 4) – which are currently represented by just a single taxon – we cannot rule out the presence of multiple independent WGDs in this group.

Finally, with respect to hybridization and polyploidy, raphid pennate diatoms have received far more attention than any other group of diatoms (see references in the Introduction and ‘Mechanisms of polyploid formation in diatoms’ sections). There is direct evidence for autopolyploid formation *in vitro* (e.g., Mann, 1994; Chepurnov and Roschin, 1995) and strong genetic evidence for natural hybrids in the few species that have been examined (Casteleyn et al., 2009; Tanaka et al., 2015). Their unique suite of traits, species richness, history of accelerated diversification, the availability established and emerging genetic model species, and extensive research on their reproductive biology establish raphid pennates as the premier lineage for uncovering the mechanisms and evolutionary consequences of polyploidy in diatoms.

### Conclusions

The phylogenomic results presented here provide strong support for a history of paleopolyploidy in diatoms that, with increased taxonomic sampling, will likely prove to be more extensive than what was uncovered with our exemplar sampling. Although WGD may be common in diatoms, its importance – if any – as a driver of speciation, lineage diversification, trait and life history evolution, or habitat shifts remains unknown. Establishing these associations, and further establishing causal links between WGD and possible evolutionary consequences, is notoriously challenging, even with the benefit of datasets much larger than those available for diatoms (Kellogg, 2016; Panchy, Lehti-Shiu, and Shiu, 2016). As with all species-rich and ecologically diverse groups, however, establishing these links represents perhaps the single greatest challenge in evolutionary research of diatoms.

Extending our sampling to more fully capture the broad ecological diversity of diatoms across environmental gradients in, for example, temperature, pH, and salinity, will help establish the evolutionary significance – if any – of WGD in diatoms. A larger comparative framework and a more precise reconstruction of the pattern and timing of paleopolyploid events – coupled with laboratory experiments – will show whether physiological shifts, either in short-term stress responses or in major habitat transitions, have been facilitated by genetic novelties introduced by gene or genome duplication. Compared to their diploid progenitors, for example, polyploid *Arabidopsis* have increased tolerance to salinity (Chao et al., 2013) – one of the principal ecological divides in diatoms and other microbial eukaryotes (Round and Sims, 1980; Mann, 1999b; Logares et al., 2009).

These results highlight numerous gaps in our understanding of diatom genomes. For a group of this size and diversity, for example, very few karyotypes and genome size estimates are available. The few data available, however, point to a level of genomic diversity and complexity that is proportional to their many other, much better known, layers of morphological and ecological diversity (Kociolek and Stoermer, 1989; Connolly et al., 2008). In light of this small but compelling collection of research, the discoveries presented herein were predictable. As genomic data for diatoms continue to accumulate, a coordinated effort to establish a reference genome dataset that captures their broad phylogenetic and ecological diversity, similar to the current call for angiosperms (Galbraith et al., 2011), is necessary to fully understand the evolution of genome size and ploidy in diatoms. Although genome size data are few, cell size data – which are available for every described diatom species – could help guide these efforts (Connolly et al., 2008). Finally, although diatoms are generally assumed to be diploid, very little is known about natural variation in ploidy levels. Few species have been surveyed for genome size data, but the intraspecific ploidy variation in two studied species, *Cocconeis placentula* (Geitler, 1973) and *Ditylum brightwellii* (Koester et al., 2010), suggests that polyploidy may play a consequential role in speciation and diversification of diatoms.

## DATA ACCESSIBILITY

RNA-seq data for XX have been deposited in the National Center for Biotechnology Information’s XX database under accessions XXX–XXX (note: GenBank submissions are in progress).

## ACKNOWLEDGEMENTS

The authors thank David Chafin, Jeff Pummill, and Pawel Wolinski for providing computational support through the Arkansas High Performance Computing Center (AHPCC), and the Chicago Botanic Garden for hosting and support of the *Treubia* and *Fabronia* computing clusters. This work was supported by the National Science Foundation (NSF) (Grant No. DEB-1353131 to AJA and DEB-1353152 to NJW), a Simons Foundation Early Career Investigator in Marine Microbial Ecology and Evolution award to AJA, and multiple awards from the Arkansas Biosciences Institute to AJA. This research used computational resources available through the AHPCC, which were funded through multiple NSF grants and/or the Arkansas Economic Development Commission, and resources available at the Chicago Botanic Garden, which were funded by NSF (DEB-1239992 and DEB-1342873 to NJW).

## APPENDICES

### Appendix 1

Sampling and assembly information for taxa used in this study.

### Appendix 2

Diatom species tree reconstruction and gene tree con-/discordance analysis. The IQ-TREE species tree reconstruction (left - phylogram, right - cladogram) of 37 diatom taxa is shown, with pie charts on cladogram indicating proportions of gene trees in congruence (blue), most common incongruence (green), all other incongruence (red) and uninformative (grey; i.e., ≤33% bootstrap support) to the shown species tree topology amongst 197 gene trees with 100% taxon occupancy and no more than 20% gaps in any alignment column. The numbers on cladogram indicate gene tree counts supporting (above) or conflicting with (below) species tree topology at a node. Support values for nodes with less than full support in IQ-TREE SH-aLRT/IQ-TREE ultrafast bootstrapping/ASTRAL/ ASTRAL-MLBS are indicated on the phylogram; * indicates a node not recovered in ASTRAL or ASTRAL-MLBS analysis.

### Appendix 3

Ks density distributions for sampled diatom taxa and *Triparma pacifica* for all Ks pipeline variations.

### Appendix 4

Results from the gene count tests of WGD across the diatom phylogeny. We used WGDgc as described in the text to estimate the birth, death and retention rate of gene duplicates assuming WGD events occurred halfway along the branch leading to the taxa specified under Hypothesis. Shown are the log likelihoods of the WGD and no WGD (null) models, the likelihood ratio (LR), degrees of freedom (df), and the probability of observing the likelihood ratio assuming a Chi-square distribution (Pchi2) with 1 degree of freedom (presence vs. absence of WGD). The taxa forming the tree used to test each putative WGD event are also shown. Species abbreviations represent the first two letters of genera and specific epithet (see also Appendix 1). *q* - retention rate, *ns* - not significant.

| Hypothesis | *q* | WGD lnL | Null lnL | LR | df | Pchi2 | SigLev | Taxa for hypothesis test |
| --- | --- | --- | --- | --- | --- | --- | --- | --- |
| gyfa | 0.148 | -33225.16 | -33515.13 | 579.936 | 1 | 0.000 | <0.001 | euna, gyfa, phtr, cram, frcy, nisp |
| asgl | 0.070 | -45831.22 | -45939.51 | 216.590 | 1 | 0.000 | <0.001 | atse, stun, euna, stsp, dite, asgl, tapo |
| tapo | 0.014 | -45933.91 | -45939.51 | 11.205 | 1 | 0.001 | <0.001 |  |
| dite | 0 | -45939.51 | -45939.51 | 0.000 | 1 | 1.000 | *ns* |  |
| asgl.tapo | 0.036 | -45915.08 | -45939.51 | 48.859 | 1 | 0.000 | <0.001 |  |
| stun | 0 | -45939.51 | -45939.51 | 0.000 | 1 | 1.000 | *ns* |  |
| atse | 0 | -45939.51 | -45939.51 | 0.000 | 1 | 1.000 | *ns* |  |
| all.but.stun | 0.011 | -64590.46 | -64596.02 | 11.126 | 1 | 0.001 | <0.001 | atse, stun, euna, gyfa, phtr, frcy, stsp, dite, asgl, tapo |
| all.but.stun.asgl.tapo | 0 | -64596.02 | -64596.02 | 0.000 | 1 | 1.000 | *ns* |  |
| thoc | 0.012 | -41506.10 | -41511.84 | 11.480 | 1 | 0.001 | <0.001 | dibr, pops, thwe, thps, skma, dips, thoc |
| thal.allo | 0 | -41511.84 | -41511.84 | 0.000 | 1 | 1.000 | *ns* |  |
| cohy | 0 | -61657.48 | -61657.48 | 0.000 | 1 | 0.998 | *ns* | cohy, leda, atse, stun, euna, odau, dibr, thps, acsu, rhse, sttu |
| leda | 0 | -61657.48 | -61657.48 | 0.000 | 1 | 0.998 | *ns* |  |
| acsu | 0 | -61657.48 | -61657.48 | 0.000 | 1 | 0.998 | *ns* |  |
| rhse | 0 | -61657.48 | -61657.48 | 0.000 | 1 | 0.998 | *ns* |  |
| acsu.rhse | 0 | -61657.48 | -61657.48 | 0.000 | 1 | 0.998 | *ns* |  |
| med.pen | 0.017 | -61643.90 | -61657.48 | 27.158 | 1 | 0.000 | <0.001 |  |
| med.pen.mel | 0.064 | -61566.87 | -61657.48 | 181.223 | 1 | 0.000 | <0.001 |  |

## FIGURE LEGENDS

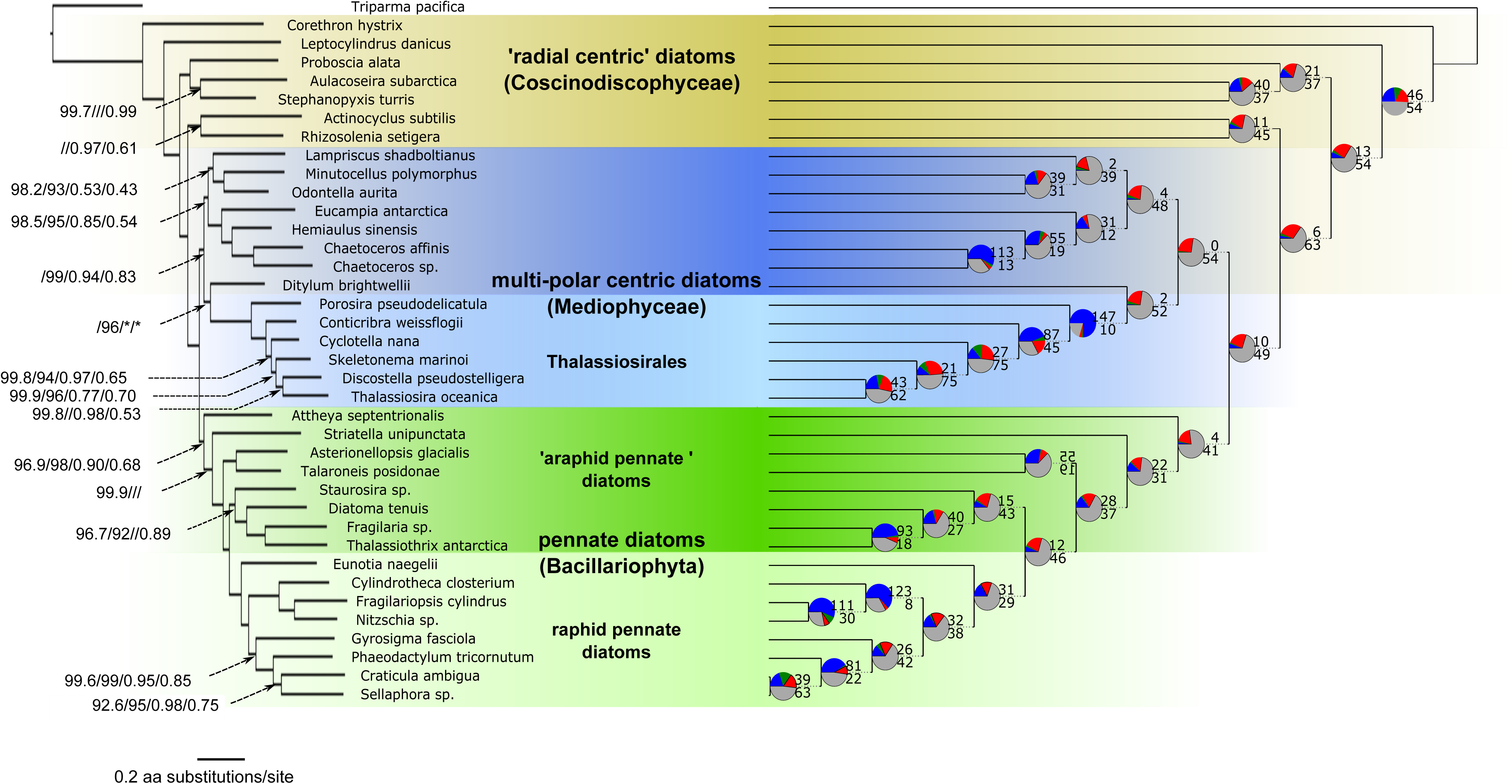

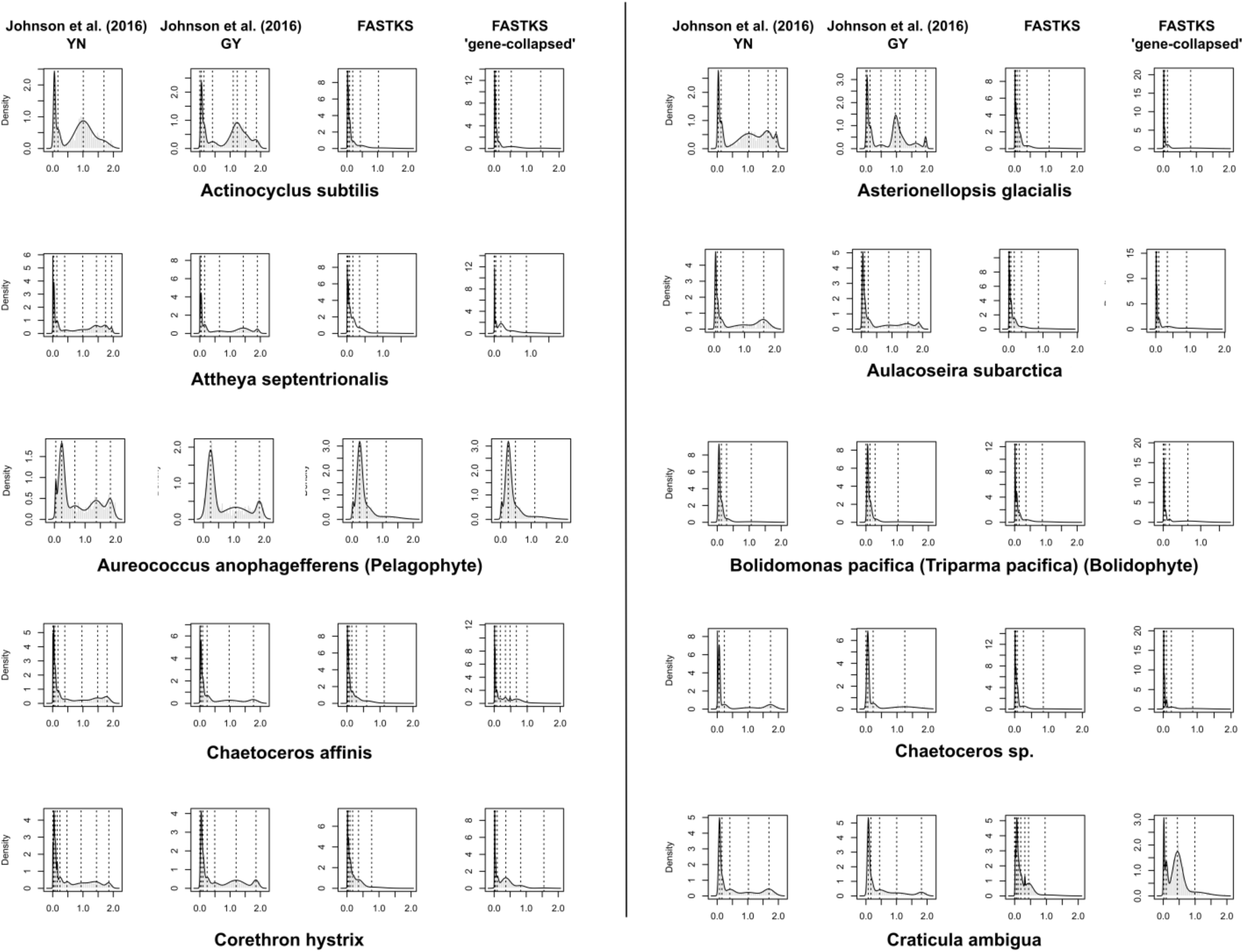

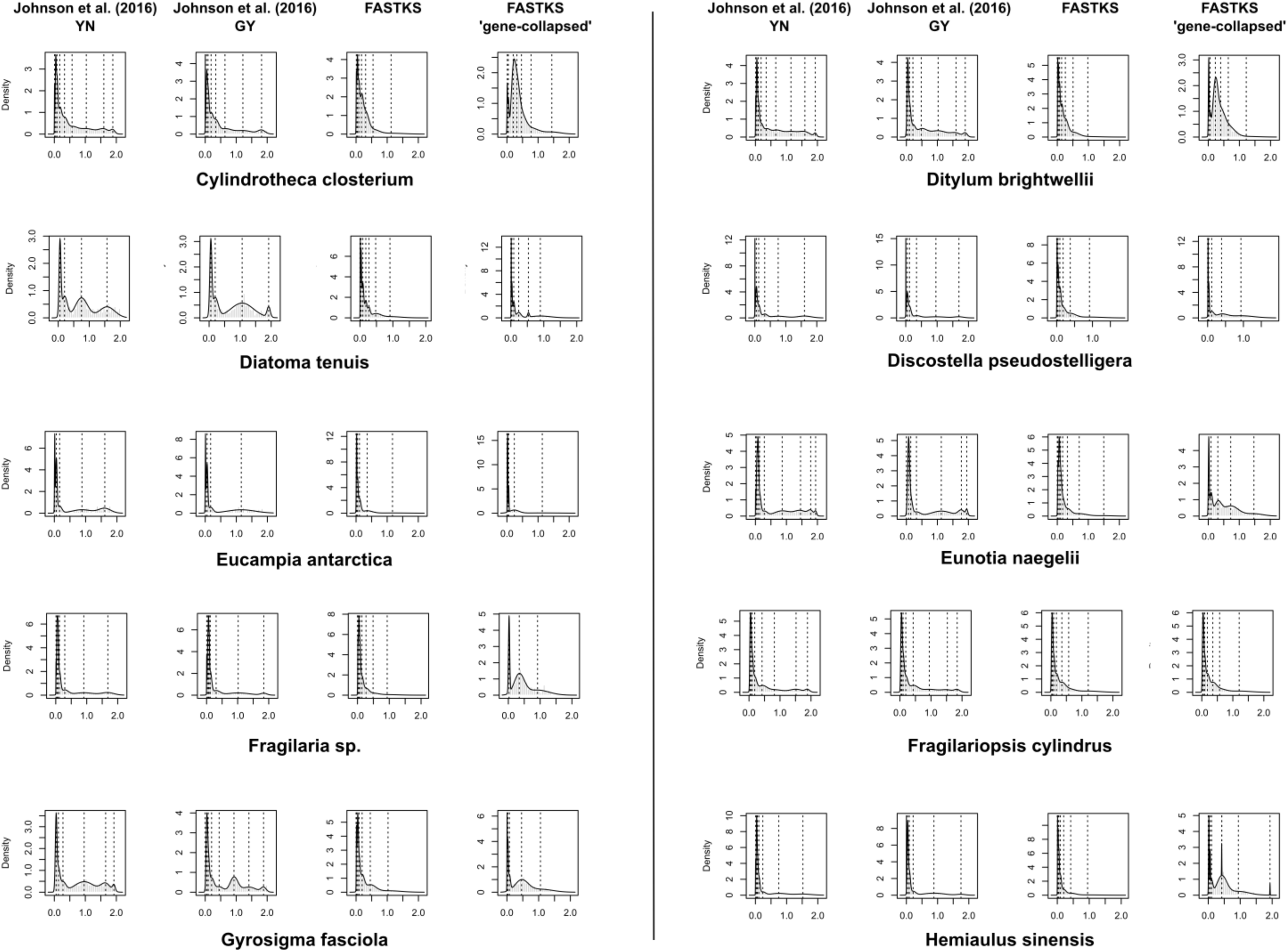

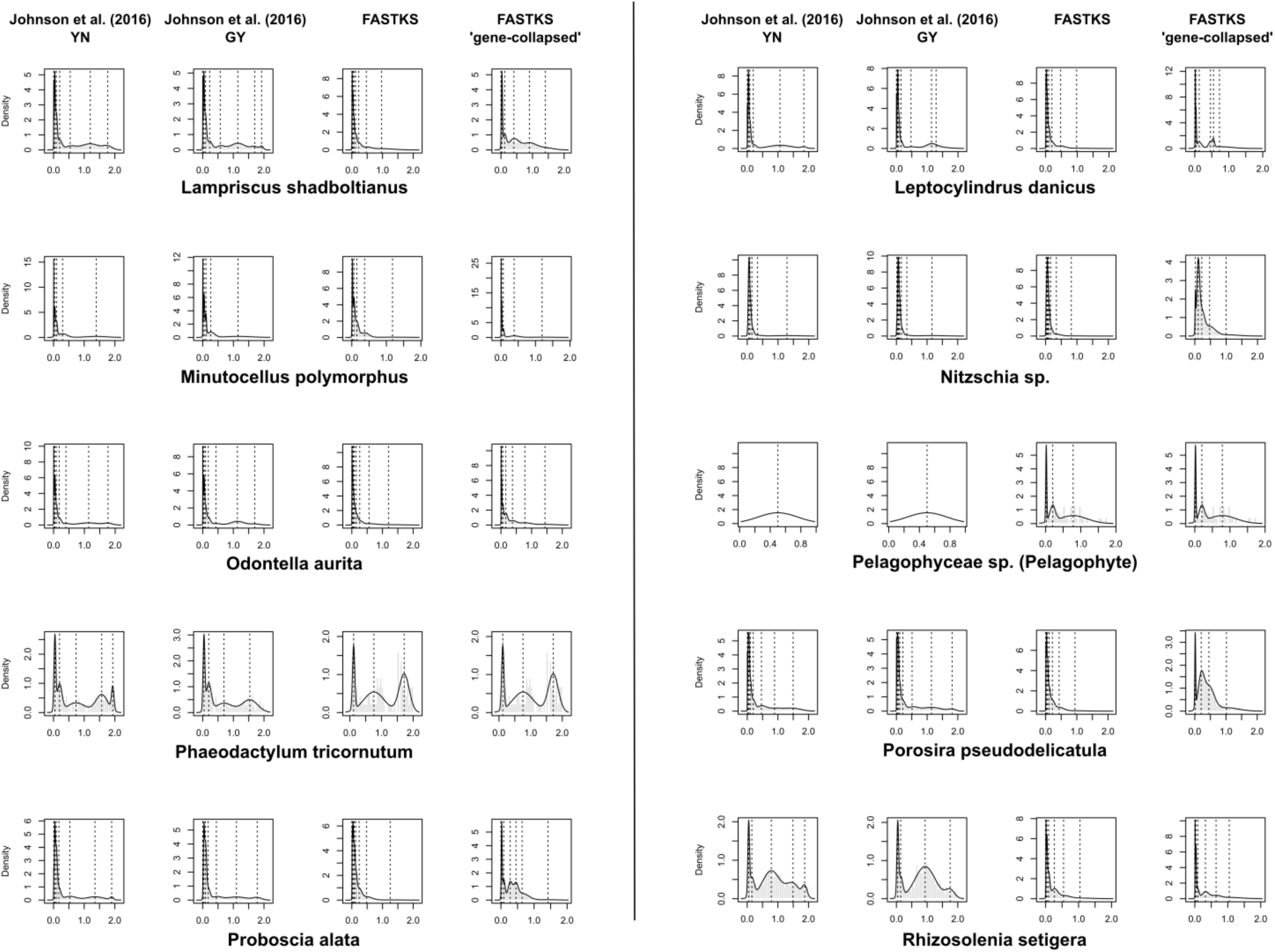

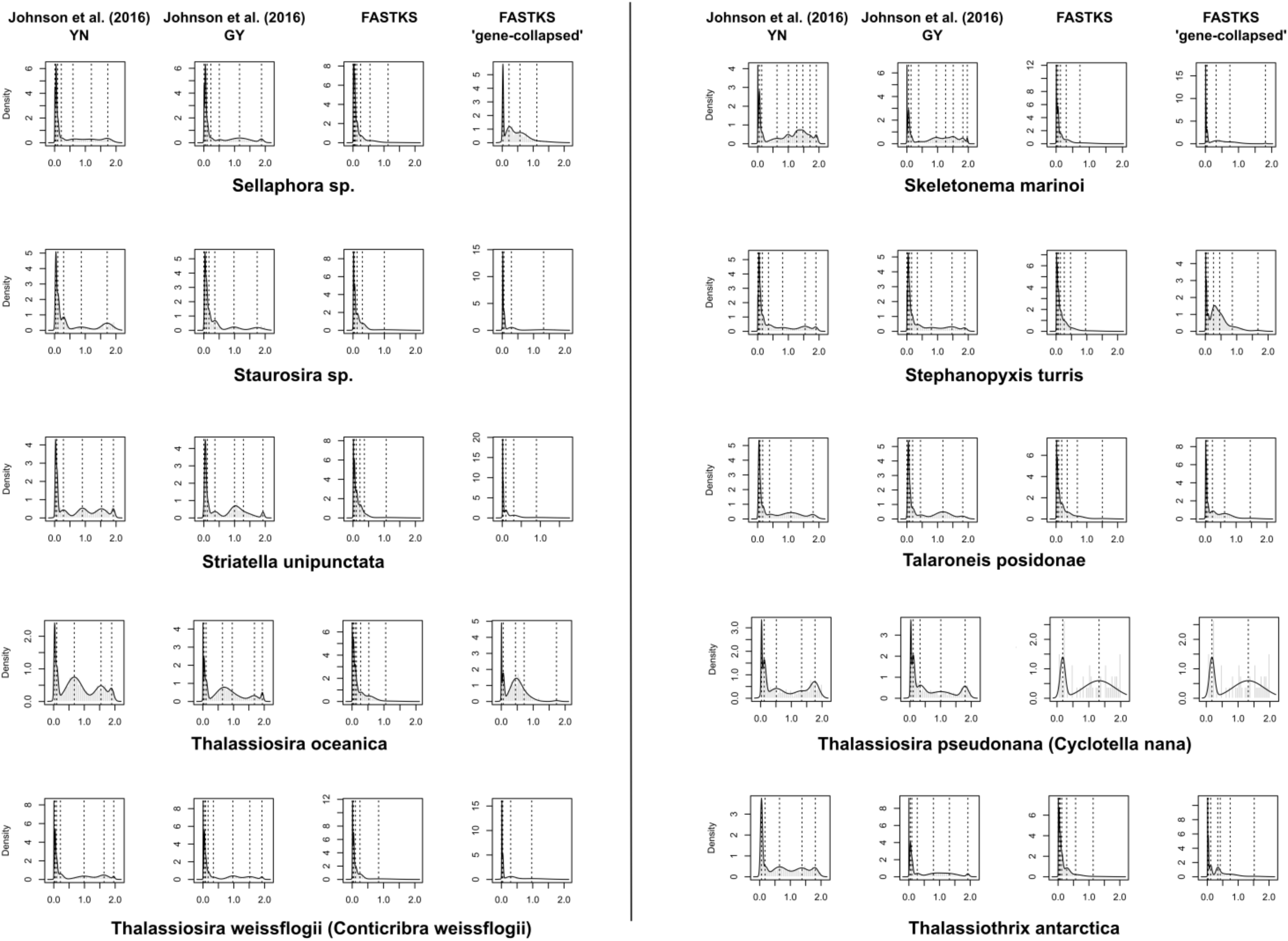

